# Integrated analysis and systematic characterization of the regulatory network for human germline development

**DOI:** 10.1101/2024.01.03.574034

**Authors:** Yashi Gu, Jiayao Chen, Ziqi Wang, Zhekai Li, Xia Xiao, Qizhe Shao, Yitian Xiao, Wenyang Liu, Sisi Xie, Yaxuan Ye, Jin Jiang, Xiaoying Xiao, Ya Yu, Min Jin, Robert Young, Lei Hou, Di Chen

## Abstract

Primordial germ cells (PGCs) are the precursors of germline that are specified at embryonic stage. Recent studies reveal that humans employ different mechanisms for PGC specification compared with model organisms such as mice. Moreover, the specific regulatory machinery is still largely unexplored, mainly due to the inaccessible nature of this complex biological process. Here, we collect and integrate multi-omics data, including 581 RNA-seq, 54 ATAC-seq, 45 ChIP-seq, and 69 single-cell RNA-seq samples from different stages of human PGC development to recapitulate the precisely controlled and stepwise process, presenting an atlas in the human PGC database (hPGCdb). With these uniformly processed data and integrated analyses, we characterize the potential key transcription factors and regulatory networks governing human germ cell fate. We validate the important roles of some of the key factors in germ cell development by CRISPRi knockdown. We also identify the soma-germline interaction network and discover the involvement of SDC2 and LAMA4 for PGC development, as well as soma-derived NOTCH2 signaling for germ cell differentiation. Taken together, we have built a database for human PGCs and demonstrate that hPGCdb enables the identification of the missing pieces of mechanisms governing germline development, including both intrinsic and extrinsic regulatory programs.

## Introduction

Human germline development is critical for fertility and newborn health. With the increasing incidences of infertility and birth defects, elucidation of germ cell development at the molecular level is critical for understanding the pathology of the related diseases^1,2^. Primordial germ cells (PGCs) are specified during early embryogenesis and ensure the establishment of the germline for subsequent generations. Due to ethical restrictions, it is impractical to access human embryos at around week 2 post-fertilization, around which human PGCs (hPGCs) are induced, to dissect the regulatory network governing hPGC specification and development^3^. To overcome this challenge, we and others established the *in vitro* strategy to induce hPGC-like cells (hPGCLCs) from pluripotent stem cells (PSCs) for characterizing the key regulatory factors for hPGC development^2,4–6^.

Human PSCs (hPSCs), including embryonic stem cells (ESCs) and induced pluripotent stem cells (iPSCs), possess the capacity to differentiate into hPGCLCs upon induction. Basically, hPSCs are induced into intermediate cell types, for example incipient mesoderm-like cells (iMeLCs) that are competent for germ cell fate induction, followed by 3-dimensional (3D) culture in aggregates for several days to induce hPGCLCs^5,7^. These hPGCLCs could be co-cultured with gonadal somatic cells to further differentiate into spermatogonia-like cells or oogonia-like cells^8,9^, establishing the *in vitro* system for uncovering the potential regulatory mechanisms governing germ cell specification and differentiation.

The development of the hPSC-derived hPGCLC induction system opened the possibility to explore the critical genes for PGC fate specification and their functions. Key transcription factors, such as SOX17, TFAP2C, PRDM1, EOMES, and PAX5, have been identified as key players for human germ cell fate determination^4,7,10–14^. These studies laid the foundation for the understanding of hPGC specification and development. However, how these transcription factors function cooperatively and how they function with other key factors remain to be further explored. Identification of more intrinsic germ cell regulators and uncovering their functions are important to understand the regulatory mechanisms for human germ cell fate. Furthermore, *in vitro* induced hPGCLCs represent nascent PGCs before the expression of DDX4 and DAZL, while co-cultured with gonadal somatic cells or injected into mouse testes promotes the differentiation of PGCLCs towards the migratory stage at which DDX4 and DAZL are expressed^5,8,9,15^. These observations suggest that signals from gonadal somatic cells are essential for the stepwise germ cell development. Identification of these soma-derived extrinsic signals is essential for understanding germ cell differentiation and reconstituting gametogenesis *in vitro*.

The development of hPGCLC induction system, together with the improvement of genomics techniques using small number of cells, lead to the accumulation of high-throughput data across different stages of germ cell development. These invaluable data provide the basis for interrogating the regulatory mechanisms governing germ cell fate. For example, integrated analysis of the scRNA-seq, ATAC-seq, and ChIP-seq data identified the TFAP2A positive progenitor cells for hPGCs and uncovered the regulation of SOX17 by TFAP2C^14^. In this study, we aimed to build a database to collect the high-throughput data of human germline cells across different developmental stages and perform integrated analyses to predict the putative transcription factors for germ cell development, uncover the regulatory network for germ cell fate, illustrate the soma-germline interaction, and explore the potential signals from somatic cells that promote the stepwise and programmed germ cell development.

## Results

### hPGC atlas in hPGCdb

In order to uncover the potential regulatory networks governing the development of human primordial germ cells (hPGCs), we aimed to establish a database collecting the data from 581 RNA-sequencing (RNA-seq), 54 Assay for Transposase-Accessible Chromatin with high-throughput sequencing (ATAC-seq), 69 single-cell RNA-seq (scRNA-seq), and 45 chromatin immunoprecipitation followed by sequencing (ChIP-seq) samples for hPGCs and related cell types (Supplementary Table 1). Largely due to the ethical restrictions, the differentiation of hPGCLCs from hPSCs has been developed and applied to define the hPGC developmental trajectory and uncover the key regulators for germ cell fate^4,5^. There are accumulating high-throughput data for hPGCs, hPGCLCs, and related cells, but lacking a database to collect the datasets from different studies to perform integrated analysis for germline regulation. Therefore, we focused on the hESCs, iMeLCs, hPGCLCs from day 1 to day 8, hPGCLCs aggregated with gonadal somatic cells termed as xenogeneic reconstituted hPGCLCs (xr-hPGCLCs), and *in vivo* hPGCs from different ages of human embryos to cover the most stages of human PGC development to reconstitute the timeline of germline development^4,7–9,11–14,16–24^ (Figure 1A and Supplementary Table 1). To minimize the variations from different analytic pipelines, we downloaded the FASTQ data and re-processed all the datasets using a unified pipeline, evaluated and screened the datasets with the same quality control criteria, and analyzed all the datasets using the standardized strategies, preparing for integrated analyses of hPGC regulation (Figure 1B). All the processed data were analyzed and visualized as interactive modules, embedded in the database website (http://43.131.248.15:6882), which is named as human primordial germ cell database (hPGCdb). hPGCdb provides 6 main functional modules including genomic tracks, heatmaps, Uniform Manifold Approximation and Projection (UMAP) plots, pseudotime curves for gene expression at single cell level, cell-cell communication, and data download^14,25–27^ (Figure S1A). In the hPGCdb main page, we provide options for visualizing RNA-seq, ATAC-seq, ChIP-seq, and scRNA-seq data based on genes and/or samples selected by users (Figure S1B). The RNA-seq data are visualized as heatmap plots (Figure 1C), the ATAC-seq and ChIP-seq data are visualized with embedded Integrative Genomics Viewer (IGV) genome browser tracks (Figure 1D-E and Figure S1C), and scRNA-seq data are visualized both in interactive two-dimension (2D) and three-dimension (3D) UMAP plots (Figure 1F-H and Figure S1D-E) after choosing samples of interest and genes/regions of interest. Customized gene expression in germline trajectory at single cell level could be presented using pseudotime curves (Figure 1I). The cell-cell interaction among different cell types generated from scRNA-seq data could be displayed after choosing samples of interest (Figure 1J). In summary, we have built a user-friendly and interactive hPGCdb based on uniformly processed high-throughput data for hPGC development.

**Figure 1.**
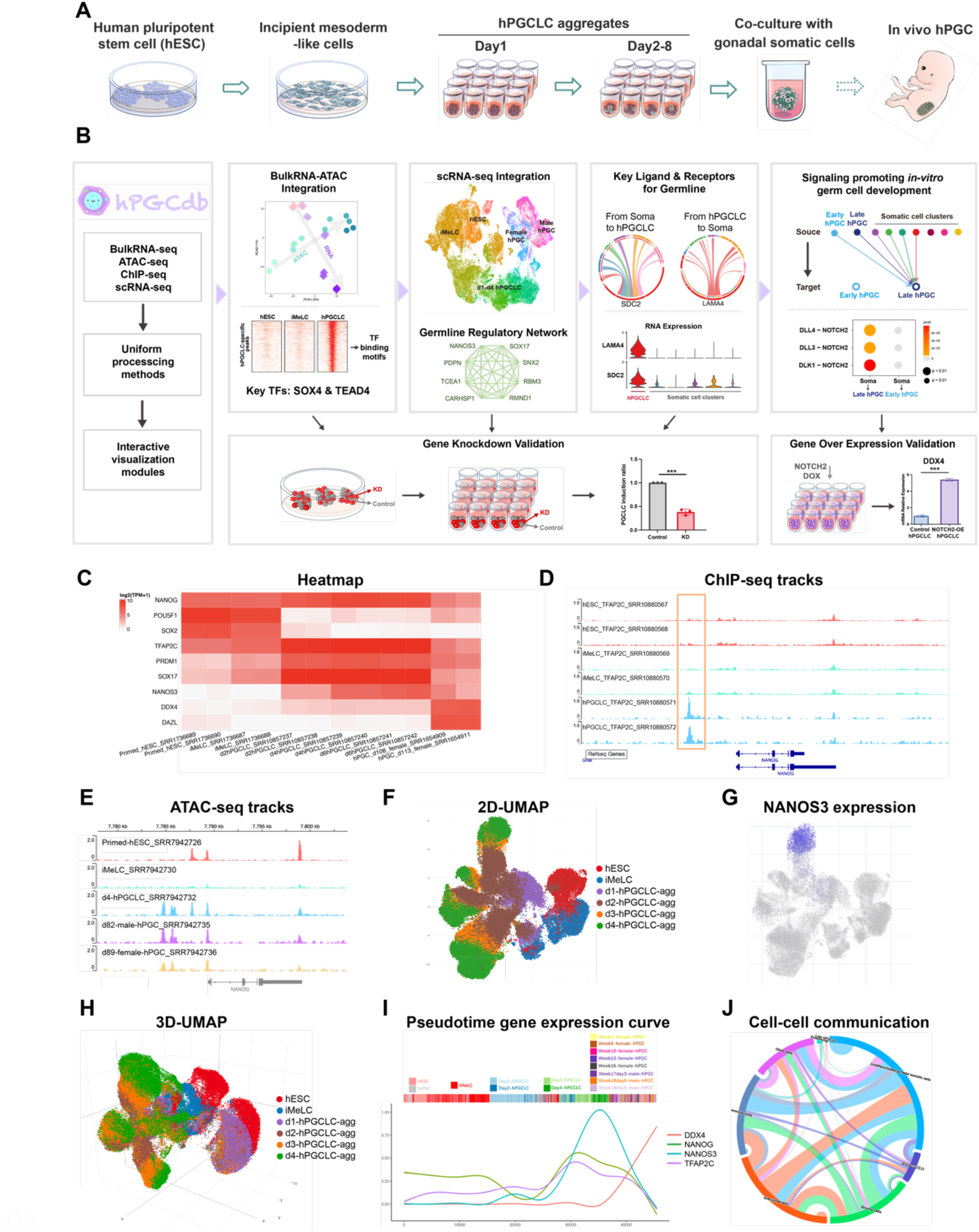
Establishment of human primordial germ cell database (hPGCdb). (A) Schematic illustration showing the cell types collected for hPGCdb. (B) Overview of study design, hPGCdb construction, integrated analyses, and experimental validation. (C) The heatmap module embedded in hPGCdb showing the expression levels of selected genes and selected samples. (D-E) The genome browser tracks embedded in hPGCdb showing the ChIP-seq (D) and ATAC-seq (E) peaks around the NANOG gene. (F-H) The 2D UMAP (F) and 3D UMAP (H) plots embedded in hPGCdb showing the cell populations of hESCs, iMeLCs, day 1-4 hPGCLC aggregates, and the expression of NANOS3 (G). (I) The pseudotime expression curve plot embedded in hPGCdb showing the time-dependent expressions of DDX4, NANOG, NANOS3, TFAP2C in the whole germline lineage cells. (J) The chord plot embedded in hPGCdb showing the cell-cell interaction of a day 70 ovary sample. Link for database: http://43.131.248.15:6882

### Identification of the transcription factors for regulating germ cell fate

After the construction of hPGCdb, we aimed to explore the transcription factors regulating hPGC development based on the integrated analysis of the multi-omics data processed and stored in hPGCdb. First, we performed principal components analysis (PCA) to integrate all the high-quality bulk RNA-seq data from different datasets after removing batch effects (Figure 1B). The samples are distributed along the differentiation trajectory from hPSCs (hESCs and hiPSCs), to iMeLCs, hPGCLCs, and *in vivo* hPGCs (Figure 2A). The germline samples express the germ cell markers at the corresponding stages (Figure S2A), indicating that these integrated samples could be applied for deeper analysis of germline regulation. We also incorporated the naïve^28^ and formative hPSCs^29^ into the PCA plot with selected samples from different stages across different datasets to further recapitulate the developmental trajectory. As expected, these samples aligned to depict the developmental trajectory along the PCA plot (Figure S2B). Based on these, we applied DESeq2^30^ to identify the differentially expressed genes (DEGs) for each developmental stage (Figure 2B). After GO term analysis we found that hPSCs (hESCs and hiPSCs) are enriched for genes related to FGF signaling; iMeLCs are enriched for genes related to gastrulation, reproductive system development, and mesoderm development; hPGCLCs are enriched for genes related to cell fate specification, gastrulation, and reproductive system development; xr-hPGCLCs are enriched for genes related to germ cell development, piRNA metabolic process, and meiosis I cell cycle process; while *in vivo* hPGCs are enriched for genes related to meiotic cell cycle, meiotic chromosome segregation, and piRNA metabolic process, respectively, aligning with each corresponding stage (Figure 2B). We also employed weighted correlation network analysis (WGCNA)^31^ to identify gene modules followed by GO term analysis and discovered the enrichment of analogous categories of biological processes (Figure S2C). Together, these analyses indicate that the data processed from different stages of germline cells in hPGCdb recapitulates the germ cell development *in vivo*, forming the basis for deep mining of the transcription factors involved in regulating germ cell fate.

**Figure 2.**
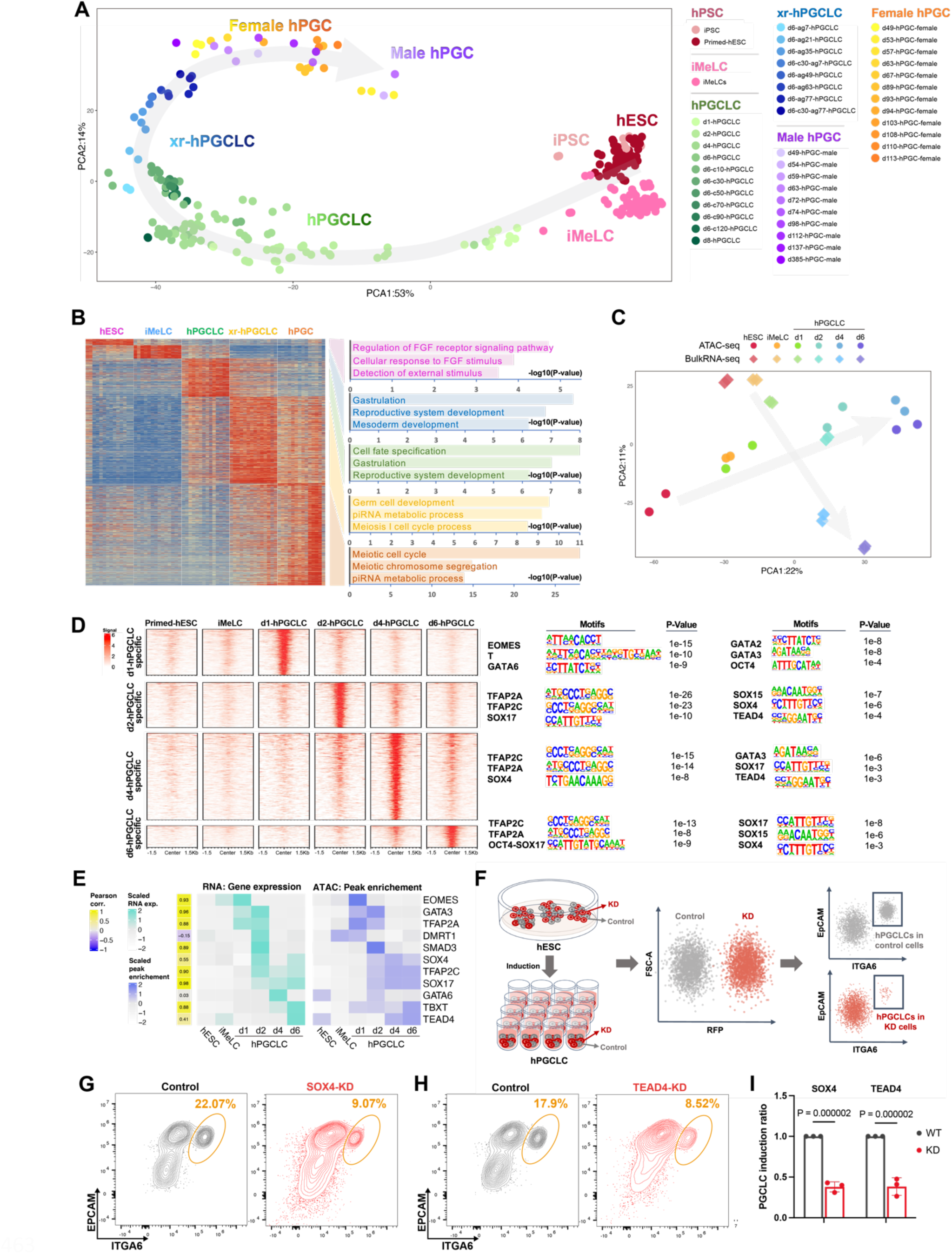
Identification of potential transcription factors important for germ cell fate. (A) PCA plot showing all of wide-type hPSCs, iMeLCs, day1-8 hPGCLCs, xr-hPGCLCs, and female and male hPGCs in hPGCdb. The curved arrow illustrates the developmental trajectory, while colors from light to deep indicate the early to late developmental stages. (B) Heatmap showing stage-specific genes (|logFC|>3) in hPSCs, iMeLCs, hPGCLCs, xr-hPGCLCs, and *in vivo* hPGCs (left panel), together with the corresponding Gene Ontology terms (right panel). (C) PCA of joint embedding of bulk RNA-seq and ATAC-seq samples. Arrows indicate the germline developmental trajectory. (D) Heatmap showing the stage-specific ATAC-seq signals in hPSCs, iMeLCs, day1 hPGCLCs, day2 hPGCLCs, day4 hPGCLCs, and day6 hPGCLCs (left), together with the corresponding transcription factor motifs enriched for these regions (right). (E) Gene expression of candidate TF regulators across cell types (middle panel). ATAC-seq peak enrichment of TF regulators across cell types (right panel). Pearson correlation coefficient between gene expression and peak enrichment (left panel). (F) Schematic illustration of the CRISPRi knockdown strategy to validate the potential role of candidate genes in hPGCLC induction. (G-H) Flow cytometry showing the induction of hPGCLCs at day 4 of aggregation differentiation using control and SOX4 knockdown hESCs (G), or control and TEAD4 knockdown hESCs (H), respectively. (I) Histogram showing the scaled proportions of hPGCLCs derived from control hESCs and gene knockdown hESCs (SOX4-KD, left; TEAD4-KD, right) as assessed by flow cytometry. The average of three replicates are shown. Error bar, standard error of the mean (SEM). The statistical significance of the differences between Control and KD are evaluated by unpaired two-sided t-test assuming unequal variances.

Next, we aimed to uncover the putative transcription factors involved in regulating different stages of germ cell development. The bulk RNA-seq samples and ATAC-seq samples from hESCs to day 6 hPGCLCs are well integrated, forming the basis for identifying key transcription factors for germ cells based on gene expression and chromatin accessibility (Figure 2C). The accessible regions unique to day1-6 hPGCLCs and hPGCs were identified, from which transcription factor motifs in the germ cell-specific open chromatin were characterized. Importantly, we discovered the known critical transcription factors, indicating the accuracy of the analytic prediction, as well as novel potential transcription factors for human germ cell development (Figure 2D, Figure S2D, and Supplementary Table 2). For example, EOMES is predicted for day 1 hPGCLCs, while SOX17 and TFAP2C are predicted for day 2-6 hPGCLCs and hPGCs, reflecting their functional stages for germ cell regulation consistent with previous studies^4,7,11,12,14^ (Figure 2D and Figure S2D). SOX4, TEAD4, and GATA family members are identified as novel potential transcription factors for regulating germ cell development (Figure 2D). To better identify the candidate regulators, we also measured regulator expression and calculated its correlation with ATAC peak enrichment across germ cell types, where SOX4 and TEAD4 show high levels for both RNA expression and chromatin accessibility at the hPGCLC or xr-hPGCLC stages. (Figure 2E and Figure S2E). Interestingly, we discovered that although the germ cell-enriched genes are not expressed in hPSCs, these gene regions are widely accessible in PSCs (Figure S3A). One example is SOX17, a critical regulators for human germ cell fate specification^4^. SOX17 is highly expressed in hPGCLCs, but not in hPSCs or iMeLCs; however, SOX17 gene region is accessible as early as at hPSC stage based on ATAC-seq analysis. This is possibly due to the binding of TFAP2C to the promoter region of SOX17 (Figure S3B). Notably, these germ cell-enriched genes are inaccessible in other embryonic somatic tissues such as embryonic liver, skin, or heart (Figure S3A), suggesting that germ cell-enriched genes are specifically accessible in embryonic cells that hold the potential to differentiate into germ cells. Meanwhile, the germ cell-specific gene modules identified from WGCNA also exhibited the same phenomenon (Figure S3C). Therefore, these observations suggest that germ cell-enriched genes are primed in embryonic stem cells for subsequent expression at later developmental stages (Figure S3D).

To validate the prediction of critical transcription factors for germ cell fate, we chose SOX4 and TEAD4 to investigate whether they play important roles in hPGCLC development by designing a fast and robust CRISPRi-based method. In this strategy, hESCs were transduced with lentiviruses bearing mCherry-labeled dCas9-CRAB for CRISPRi and the corresponding gRNAs. We controlled the titers of the lentiviruses to achieve that about half of the hESCs were RFP (mCherry) positive (knock-down, KD), while the other half were RFP negative (control cells). The mixture of RFP positive and RFP negative cells was used to induce hPGCLCs, followed by flow cytometry analysis at day4 for hPGCLC induction efficiency using ITGA6 and EPCAM as surface markers (Figure 2F). The advantage of this strategy is that both KD and control cells are induced into hPGCLCs in the same wells, minimizing the batch variations. In control hESCs, distinct hPGCLC population was shown using ITGA6 and EPCAM as surface markers for flow cytometry. Notably, the putative hPGCLC population was dramatically decreased (from 22.1% to 9.1%) when SOX4 was knocked down (Figure 2G, 2I, and Figure S3E). Similarly, the hPGCLC induction was also severely impaired (from 17.9% to 8.5%) when TEAD4 was knocked down (Figure 2H, 2I, and Figure S3E). These results indicate that SOX4 and TEAD4 are important for hPGCLC induction or development. To further examine whether the identified potential transcription factors are related to germline diseases, we determined the percentage of the identified transcription factors assigned to various reproduction-related diseases. Importantly, the identified transcription factors exhibit significant correlations with 13 types of infertility-related diseases and germ cell tumors compared to randomly selected genes (Figure S3F). Taken together, the data collected and processed in hPGCdb could be applied to identify the potential key transcription factors for human germline development, based on which we discovered that SOX4 and TEAD4 are novel transcription factors for regulating human germ cell fate.

### Elucidation of the regulatory networks for human germ cell development

To further characterize the potential regulators for germ cell development, we collected scRNA-seq data of hESCs/hiPSCs, iMeLCs, hPGCLCs, xr-hPGCLCs, and *in vivo* male PGCs (day122-131 fetal testes) and female PGCs (day49-91 fetal ovaries) across different developmental stages from the four datasets in hPGCdb to construct the germline trajectory at single cell level (Figure1B). First, for the three datasets covering hPSCs to hPGCLCs, we applied the time-constraint URD (tcURD)^14,26^ to identify differentiation trajectories unbiasedly by building up the single cell differentiation trees (Figure 3A, 3C, 3E). Then the germline trajectory segment was determined based on the expression of the germ cell specific gene NANOS3 or DDX4 (Figure 3B, 3D, 3F). For the scRNA-seq data from fetal gonads, we extracted the germ cells based on the expression of PGC markers such as NANOS3, TFAP2C, DDX4, and DAZL, because there is no PGC progenitors or earlier cells at these stages (Figure S4A-C). Next, we integrated all cells from the germline trajectory together after removing the batch effects by Mutual Nearest Neighbors (MNN)^32^. The cells embedded using UMAP aligned with the germline development from hPSCs to iMeLCs, day1-day4 hPGCLCs, d49-d131 female and male hPGCs, indicating the validity of scRNA-seq data integration (Figure 3G-I). Based on these integrated germline cells, we performed the single cell WGCNA (scWGCNA)^33^ to explore the potential regulatory modules governing human germ cell development. Two regulatory modules drew our attention because they both reflected the potential regulation at the hPGCLC stage (Figure 3J-M). Notably, known genes important for hPGCLCs were identified at these two modules, such as SOX17, NANOS3, and PRDM1 (Figure 3K, M, Supplementary Table 2). To further detect the expression patterns of the hub genes in these two modules, we examined the expression levels of these hub genes using the bulk RNA-seq data collected in Figure 2. Consistently, most of these genes are enriched in hPGCLCs (Figure 3N and Figure S4E), suggesting that they may play critical roles for regulating germ cell development. We also identified a regulatory module that may play critical roles in TFAP2A+ progenitors for hPGCLCs^14^, given the expression of TFAP2A, GATA2, GATA3, and CDX2 (Figure S4F-G, Supplementary Table 2). To test whether we may predict the germline regulatory network without removing somatic cells, we applied scWGCNA to the UCLA2 dataset containing all cells from hESCs, iMeLCs, hPGCLC aggregates day 1 to day 4 (Figure S4H). Notably, we identified a regulatory gene module that contains the hub genes TFAP2C and SOX17, which are the key regulators for hPGCLC specification (Figure S4I-K).

**Figure 3.**
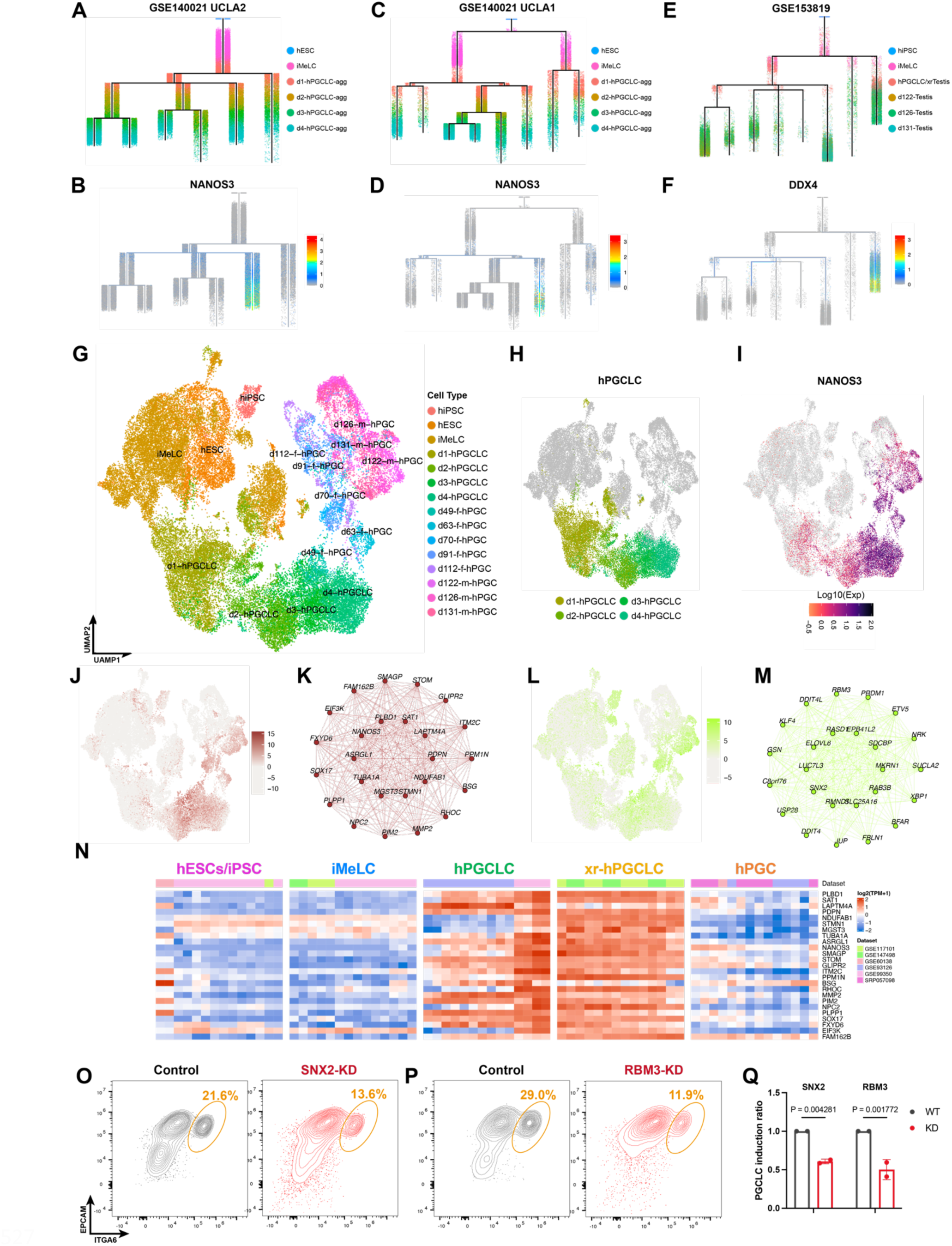
Characterization of potential regulatory networks governing human germ cell development. (A-F) URD trees showing the developmental trajectories established from the three datasets: GSE140021-UCLA2 (A), GSE140021-UCLA1 (C), GSE153819 (E). The expression of NANOS3 indicates the establishment of the early hPGCLCs in UCLA2 (B) and UCLA1 (D), while the expression of DDX4 represents the establishment of the late hPGCs in GSE153819 (F), contributing to identify and extract the single cells in the germline lineage from each dataset. (G) UMAP plot showing the integrated germline lineage cells containing cells at all stages involved in germ cell development after removing batch effect. (H) day1-4 hPGCLCs are highlighted in the UMAP plot of the integrated germline lineage cells. (I) The expression of NANOS3 in the UMAP plot of integrated germline lineage cells. (J-M) Two putative regulatory network modules for early PGCs based on scWGCNA (K, M), and the corresponding expression patterns of the two modules (J, L). (N) The expression of hub genes identified from the germ cell regulatory network modules in (K). (O) Flow cytometry showing the induction of hPGCLCs at day 4 of aggregation differentiation using control and SNX2 knockdown hESCs (O), or control and RBM3 knockdown hESCs (P), respectively. (Q) Histogram showing the scaled proportions of hPGCLCs derived from control hESCs and knockdown hESCs (SNX2-KD, left; RBM3, right) as assessed by flow cytometry. The average of two replicates are shown. Error bar, standard error of the mean (SEM). The statistical significance of the differences between control and KD are evaluated by unpaired two-sided t-test assuming unequal variances.

To test whether the identified hub genes are important for hPGCLC induction, we applied the CRISPRi strategy we established (Figure 2F) to the hub genes related to post-transcriptional regulation or developmental diseases. We first chose SNX2 (related to Parkinson Disease and Late-Onset), RBM3 (a member of the glycine-rich RNA-bindinf protein family), and RMND1 (related to Mitochondrial Oxidative Phosphorylation Disorder) from the human germline lineage WGCNA networks (Figure 3K and 3M). Importantly, CRISPRi knockdown of SNX2, RBM3, or RMND1 led to the significant reduction of hPGCLC percentage (Figure 3O-Q, Figure S5A and S5D-E), suggesting that these genes are involved in the regulation of hPGCLC induction. We also chose CARHSP1 (an RNA-binding protein regulating the target mRNA stability) and TCEA1 (associated to translation elongation factor activity) from the WGCNA hub genes identified in germ cells with somatic cells (Figure S4J). Importantly, the hPGCLC population was significantly decreased in the knockdown groups compared with control groups (Figure S5B-E), supporting that the identified regulatory gene modules may play important roles for human germ cell development. Altogether, these results indicate that the scRNA-seq data collected in the hPGCdb could be applied to uncover the potential key regulatory networks for human germline development, from which we uncovered several gene modules as important germline regulators.

### Characterization of the germline-soma interaction during hPGCLC development

Germ cell development is accompanied with changing niches during gametogenesis^34^. To characterize the potential regulatory roles from surrounding somatic cells, we calculated the cell-cell interaction between UCLA2 day 4 hPGCLCs and somatic cells using CellChat^27^ (Figure 1B). Based on the expression of NANOS3 and other markers for different germ layers, we annotated the 6 populations in day 4 hPGCLC aggregates (Figure 4A-B and Figure S6A). Rich interaction between somatic cells and hPGCLCs was identified (Figure 4C-E and Figure S6B), suggesting that hPGCLC development requires the interaction with surrounding somatic cells. Among all the interaction pairings, the hPGCLC surface marker ITGA6 was found to be enriched in hPGCLCs as receiving cells (Figure 4C), validating our cell-cell interaction analysis. Importantly, Syndcan-2 (SDC2) in hPGCLCs as receiving cells and Laminin Subunit Alpha 4 (LAMA4) in hPGCLCs as sending cells drew our attention (Figure4C-F), as Syndcan families have been discovered to be involved in testicular germ cell tumors and in regulating germ cells via Notch signaling^35,36^, while Laminin has been implicated in PGC migration in Drosophila and mice^37,38^. We then examined the expression levels of SDC2, LAMA4, and ITGA6, confirming their specific expression pattern in hPGCLCs (Figure 4G, 4J, and S6C).

**Figure 4.**
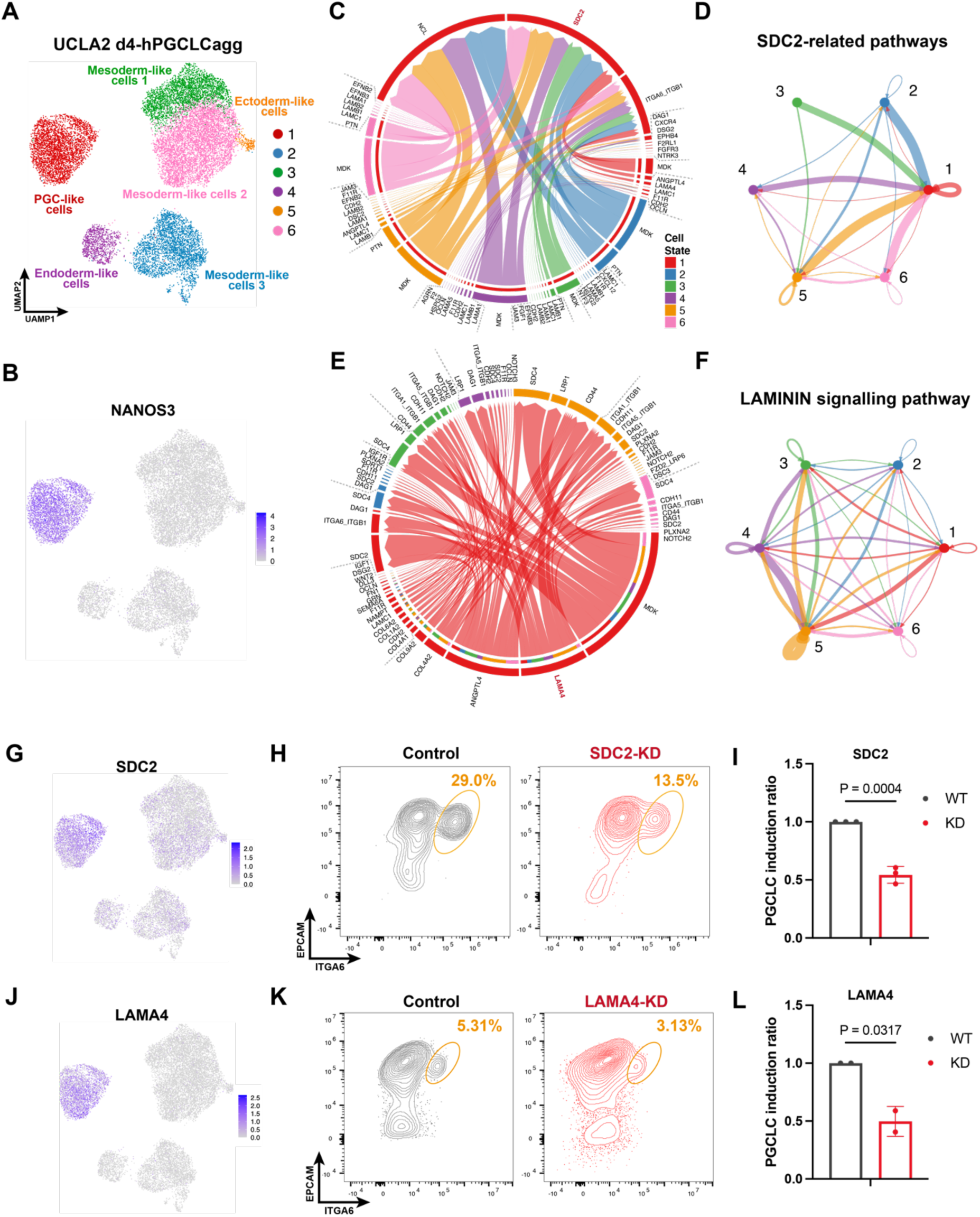
Elucidation of the soma-germline interaction in hPGCLC aggregates. (A) UMAP plot showing the cell clusters of UCLA2 day 4 hPGCLC aggregate samples. (B) UMAP plot showing the expression of NANOS3 in UCLA2 day 4 hPGCLC aggregate samples. (C) Chord plot showing the cell-cell interactions targeting to day4 hPGCLCs from various somatic cell clusters. (D) SDC2-related cell-cell interactions among all cell clusters in UCLA2 day 4 hPGCLC aggregates. (E) Chord plot shows cell-cell interactions sending from day4 hPGCLC to various somatic cell clusters. (F) LAMININ signaling-related interactions among all cell clusters in UCLA2 day 4 hPGCLC aggregates. (G) UMAP plot showing the expression of SDC2 in UCLA2 day 4 hPGCLC aggregate samples. (H) Flow cytometry showing the induction of hPGCLCs at day 4 of aggregation differentiation using control and SDC2-KD hESCs. (I) Histogram showing the scaled proportions of hPGCLCs derived from control hESCs and SDC2 knockdown hESCs as assessed by flow cytometry. The average of two replicates are shown. Error bar, standard error of the mean (SEM). The statistical significance of the differences between Control and KD are evaluated by unpaired two-sided t-test assuming unequal variances. (J) UMAP plot showing the expression of LAMA4 in UCLA2 day 4 hPGCLC aggregate samples. (K) Flow cytometry showing the induction of hPGCLCs at day 4 of aggregation differentiation using control and LAMA4-KD hESCs. (L) Histogram showing the scaled proportions of hPGCLCs derived from control hESCs and LAMA4-KD hESCs as assessed by flow cytometry. The average of two replicates are shown. Error bar, standard error of the mean (SEM). The statistical significance of the differences between control and KD are evaluated by unpaired two-sided t-test assuming unequal variances.

To confirm the identified soma-germline interaction, we next performed cell-cell interaction analysis using UCLA1 scRNA-seq data containing day 4 hPGCLCs and surrounding somatic cells (Figure S7A-B), which also revealed rich soma-germline interaction (Figure S7C). The main roles of SDC2 and LAMA4 in soma-hPGCLC interaction (Figure S7D-G) and their specific expression in hPGCLCs (Figure S7H-J) also inferred their potential roles. These observations are consistent with the findings in UCLA2 and promoted us to test whether SDC2 and LAMA4 are important for hPGCLC development. To test this, we applied the CRISPRi strategy we established to knock down SDC2 and LAMA4, and discovered that hPGCLC population was significantly decreased upon SDC2 knockdown (Figure 4H-I and Figure S7K) or LAMA4 knockdown (Figure 4K-L and Figure S7K) compared with control, respectively, suggesting that SDC2-mediated and LAMA4-mediated soma-germline interaction is important for hPGCLC differentiation. Taken together, the scRNA-seq datasets in the hPGCdb could be applied to dissect the cell-cell interaction between germ cells and somatic cells, from which we discovered that SDC2 and LAMA4 may play important roles for mediating soma-germline interaction to regulate germ cell development.

### NOTCH2 regulates the proceeding of germline development

To uncover the potential mechanisms for regulating progressive development of germline, we further explored the soma-germline interaction by concentrating on *in vivo* fetal gonads containing both early hPGCs at nascent stage and late hPGCs at around migratory stage (Figure 1B). One long-standing and critical question for human germ cell development is how the PGCs proceed from early (DDX4-negative) to late (DDX4-positive) stage. For hPGCLCs from 3D induction system, they represent the pre-migratory stage before DDX4 is expressed^3,39^. Importantly, in the non-human primate, when day 4-6 PGCLCs were transplanted into the mouse testicles after germ cells were depleted by radiation, these DDX4-negative germ cells differentiated into the DDX4-positive cells by receiving the signals from the gonadal somatic cells of recipient mice^15^. This observation indicates that signals from gonadal somatic cells are critical for promoting hPGC differentiation towards subsequent stages. To explore the potential signaling pathways that regulate the transition from DDX4-negative to DDX4-positive stages, we focused on the scRNA-seq data from day 70 fetal ovaries, which contain both DDX4-negative and DDX4-positive PGCs (Figure 5A). The DDX4 negative PGCs express NANOS3, TFAP2C, SOX17, and TEAD4, supporting that they are early stage PGCs (Figure 5B). Conversely, the DDX4 positive PGCs express late PGC markers, including DAZL, MAEL, and SYCP3 (Figure 5C). We also determined the somatic cell fates in this sample, including undifferentiated gonadal somatic cells (LHX9 positive), stromal cells (TCF21 positive), endothelial cells (PECAM1 positive), granulosa cells (WNT6 positive), and immune cells (PTPRC positive)^40^ (Figure S8B-C). Based on the annotated cell groups above, we could compare the soma-germline interaction for these two PGC populations (DDX4 negative population versus DDX4 positive population) to identify the potential signaling pathways that trigger the differentiation of PGCs from DDX4 negative stage to DDX4 positive stage.

**Figure 5.**
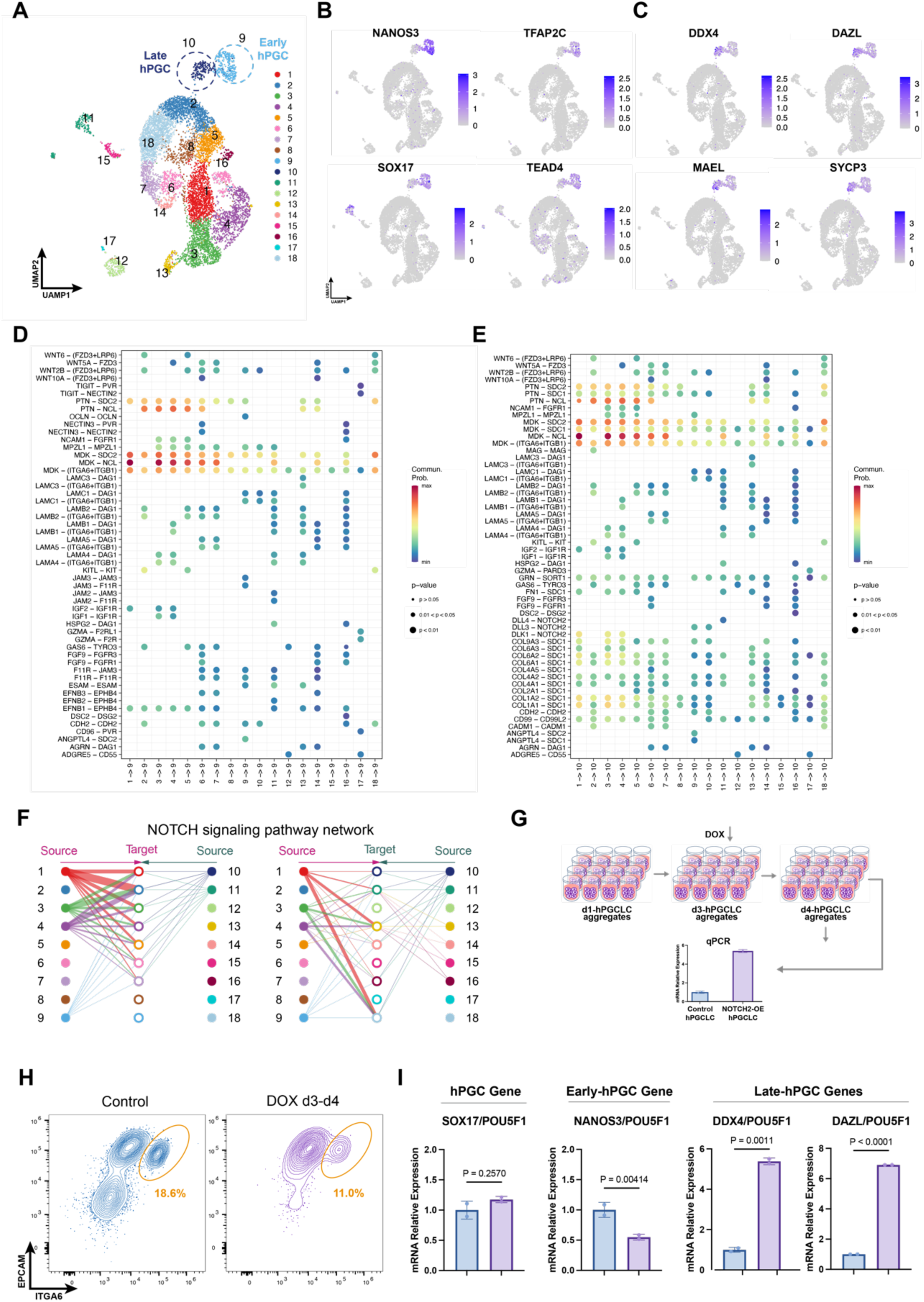
NOTCH2 signaling may regulate the transition from DDX4 negative to DDX4 positive hPGCLCs. (A) UMAP plot showing all cells from day70 ovaries, where the early hPGC cluster (cluster 9) and the late hPGC cluster (cluster 10) are circled, respectively. (B-C) UMAP plots showing the expression levels of the early PGC marker gene (NANOS3, TFAP2C, SOX17, TEAD4) and the late PGC marker gene (DDX4, DAZL, MAEL, SYCP3) in the day70 ovaries, contributing to the annotation of the early and the late hPGC clusters, respectively. (D-E) Dot plot showing the putative ligand-receptor pairing sending from various somatic cell clusters to the early hPGC cluster (D) and sending from various somatic cell clusters to the late hPGC cluster (E). (F) The communication network of NOTCH2 signaling pathway. Cluster 9 and cluster 10 are the early hPGC cluster and the late hPGC cluster, respectively. (G) Schematic illustration of the experimental design of NOTCH2 overexpression in hPGCLCs. (H) Flow cytometry showing the induction of hPGCLCs at day 4 of aggregation differentiation using control (left) and NOTCH2 over-expression hESCs (right). (I) Histogram showing expression levels of common hPGC genes, early hPGC marker gene, and late hPGC marker genes in control hESCs and NOTCH2-OE hESCs. The average of two replicates are shown. Error bar, standard error of the mean (SEM). The statistical significance of the differences between control and KD are evaluated by unpaired two-sided t-test assuming unequal variances.

We applied CellChat to characterize the soma-germline interaction for DDX4 negative and DDX4 positive stages, respectively (Figure 5D-E, Figure S8D, Figure S9). We hypothesized that signaling pathways active exclusively in DDX4 positive PGCs but not in DDX4 negative PGCs are crucial for promoting the differentiation from early stage to late stage PGCs. Interestingly, various somatic cell clusters send NOTCH2 signaling only to DDX4 positive stage but not DDX4 negative stage (Figure 5D-F), which is consistent with previous findings^39,40^. To test whether NOTCH2 is sufficient to promote the differentiation of PGCs from DDX4 negative stage to DDX4 positive stage, we ectopically activated NOTCH2 signaling during hPGCLC development. We reasoned that NOTCH2 signaling should be inactive during early stage of hPGCLC development, but turn to be active to promote the differentiation of hPGCLCs towards late stage. Therefore, we added Dox to over-express the activated form of NOTCH2 (intracellular domain of NOTCH2) at day3, and harvested the day 4 hPGCLCs for analysis (Figure 5G). As expected, over-expression of NOTCH2 decreased the hPGCLC population at day 4 (Figure 5H). But importantly, over-expression of NOTCH2 leads to the significant up-regulation of late stage PGC markers DDX4 and DAZL, down-regulation of early stage PGC maker NANOS3, and comparable expression of SOX17 (Figure 5I and Figure S8A), suggesting that NOTCH2 signaling is involved in promoting the differentiation of hPGCs from DDX4 negative stage to DDX4 positive stage. Taken together, these results indicate that the scRNA-seq data collected in the hPGCdb could be applied to investigate the soma-germline interaction and the potential critical regulatory signaling pathways governing the precise and stepwise gametogenesis.

## DISCUSSION

Specification of PGCs during early embryogenesis is essential for the establishment of germline for propagating genetic and epigenetic information from parents to offspring. In this study, we have built a database for integrated analysis of the regulation of human germline specification and differentiation by collecting high-throughput multi-omics data representing the precise and stepwise development of the human germ cells. Combining RNA-seq and ATAC-seq, we identified the potential critical transcription factors for germ cell fate. Based on scRNA-seq analysis, we systematically characterized the putative regulatory network and soma-germline interaction that govern the human germ cell development.

Human germline development is initiated at around day 12-14 after fertilization with the specification of primordial germ cells^14,41^. Due to the ethical restrictions and limited sources of human embryos, it is impractical to investigate human embryos directly for germline specification and development. Therefore, mice have been served as the model system to understand mammalian PGCs for decades^3,42,43^. However, with the establishment of the *in vitro* system that induces PGCLCs from PSCs, more and more studies emphasize the divergence of germline specification between humans and mice. For example, SOX2 is important for mouse PGCs but dispensable for human germline specification^44^, while SOX17 is critical for human PGC induction^4^, although not necessary for mouse PGCs^45^. PRDM14 is one of the essential determinants for mouse PGC fate but plays different roles in human PGCs^23,46,47^. Furthermore, even the origin of PGCs is probably different in humans and mice. Mouse PGCs derive from posterior epiblasts while human and non-human primate PGCs may be specified from amnion progenitors^3,43,48^. Altogether, it is important to investigate human germline specification and development using the *in vitro* hPGCLC system together with the *in vivo* embryonic hPGCs. Therefore, we built the hPGCdb for collecting the high-throughput data of both *in vitro* and *in vivo* human germ cells from different developmental stages for interrogating the potential regulatory mechanisms governing human germ cell development.

Our integrated analysis not only identified the known transcription factors essential for human germ cell fate including EOMES, TFAP2C, SOX17, and TFAP2A, but also novel transcription factors such as SOX4 and TEAD4. CRISPRi knockdown of SOX4 or TEAD4 leads to dramatic decrease of hPGCLC induction, confirming that they are important for human germline development (Figure 2). SOX4 is involved in the regulation of stemness and differentiation, and regulates developmental processes through pathways including PI3K, WNT, and TGFβ signaling^49^. SOX4 also participates in the reactivation of embryonic gene expression program during wound healing in mice^50^. Interestingly, SOX4 is mainly expressed in the gonadal somatic cells in mice to regulate gonadal morphogenesis and germ cell differentiation^51^. However, SOX4 is expressed in hESCs/hiPSCs, iMeLCs, and hPGCLCs, and involved in the regulation of hPGCLC induction (Figure 2D, 2E, 2G, and Figure S2E), which may further imply the divergence of germline development in humans and mice. The detailed functions of SOX4 and molecular mechanisms through which SOX4 regulates the development of hPGCLCs warrant further investigation. TEAD4, as the critical cell fate determinant of trophectoderm^52–54^, is also expressed in hPGCLCs and involved in hPGCLC induction (Figure 2D, 2E, 2H, and Figure S2E). The functions of TEAD4 in germline development remain mostly unknown. Our integrated analysis has also revealed the potential networks for the regulation of human germ cell development (Figure 3 and Supplementary Table 2). All these candidate intrinsic germline regulators could be applied to CRISPR-KO or CRISPRi screening to get a comprehensive understanding of the regulation of human germ cell specification and development.

We also explored the extrinsic regulation of human germline development by investigating soma-germline interaction during hPGCLC and hPGC development. Mouse and human PGCs are specified by the signals from extraembryonic tissues, such as BMP4^3^. PGCs interact with somatic cells after specification, during migration and colonization with gonadal somatic cells^34^. In this study, we have identified SDC2 and LAMA4 as potential soma-germline interacting factors that regulate hPGCLC induction (Figure 4, S5 and S6), suggesting that the soma-germline interaction could be recapitulated in hPGCLC aggregates. One key bottleneck for hPGCLC differentiation is from nascent PGC stage (NANOS3 positive, DDX4 negative) to migrating PGC stage (NANOS3 negative, DDX4 positive). Notably, these *in vitro* PGCLCs could reach the DDX4 positive stage upon transplantation into the mouse testicle^15^, implying that signals from gonadal somatic cells promote the proceeding of germ cell development. In the scRNA-seq data of a day 70 embryonic gonad sample in the hPGCdb, both nascent PGCs (NANOS3 positive, DDX4 negative) and migrating PGCs (NANOS3 negative, DDX4 positive) are present (Figure 5A-C). Comparison of the soma-germline interactions identified the NOTCH2 signaling as a candidate pathway that regulates the differentiation for hPGCLC from nascent stage to migrating stage. This was verified by the over-expression of the activated form of NOTCH2 that triggered the up-regulation of the DDX4 and DAZL, markers for migrating PGCs (Figure 5H-I and Figure S8A). These results emphasize the importance of niche cells in regulating the development of germ cells. However, how NOTCH2 signaling activates the expression of DDX4 and DAZL for promoting PGC differentiation requires further investigation. One recent study identified DMRT1 as the key factor that triggers the expression of DAZL but not DDX4^55^. Whether DMRT1 is the downstream effector of NOTCH2 is unknown. Nevertheless, we discovered that activation of NOTCH2 signaling in hPGCLCs triggers the expression of both DAZL and DDX4, forming the basis to improve the *in vitro* differentiation system towards functional gametes for regenerative medicine, as well as to facilitate the dissection of molecular mechanisms ensuring the precise and stepwise differentiation of germ cell lineage.

In summary, we have built the hPGCdb for systematic analysis of both intrinsic and extrinsic regulation of human germline development. These invaluable resources will contribute to the understanding of human germ cell fate specification and the tightly controlled stepwise differentiation of human PGCs, which will also provide potential pathological mechanisms for fertility-related disorders and diseases.

## FIGURES AND LEGENDS

**Figure S1.**
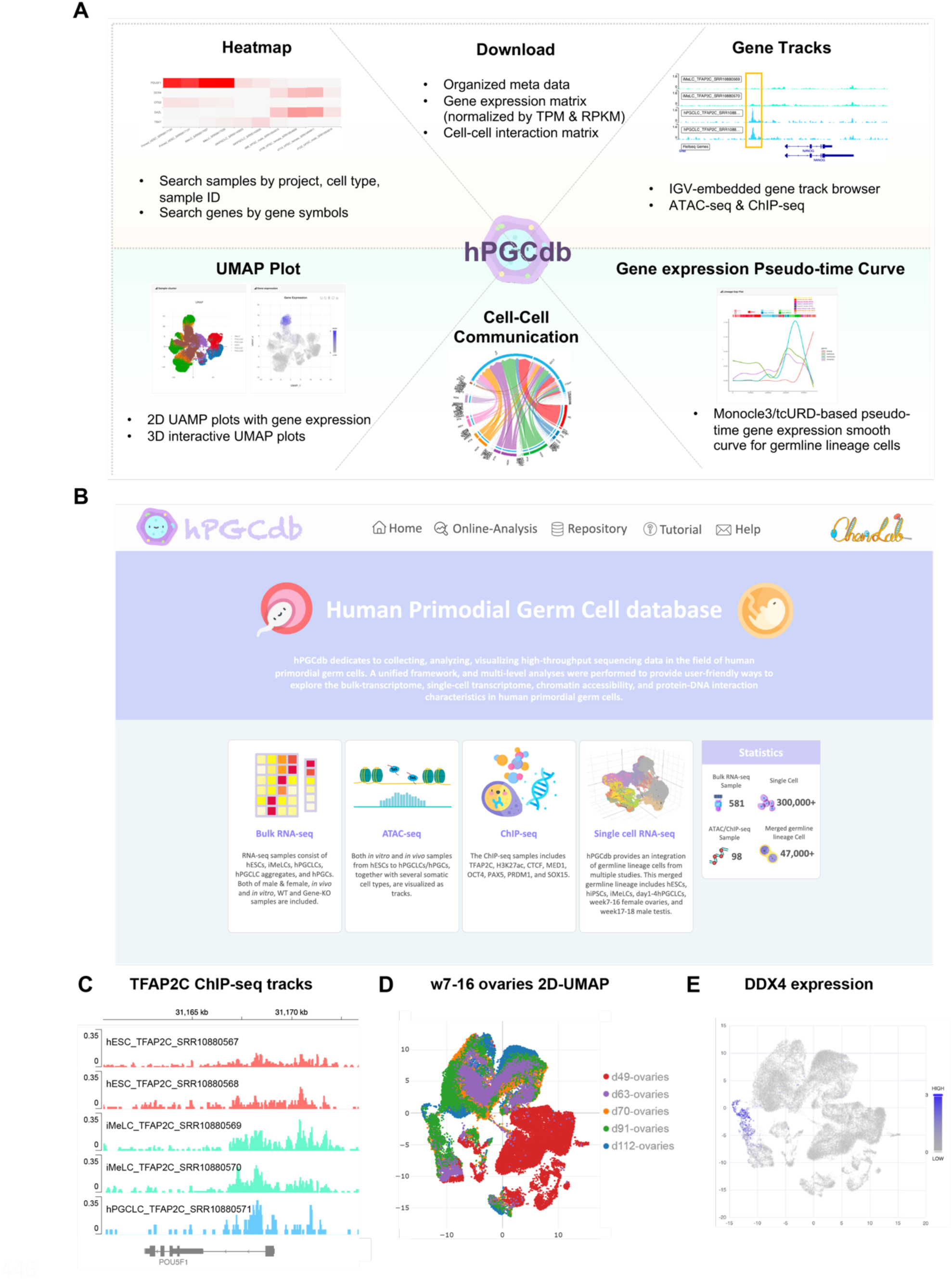
hPGCdb database for visualization of germline cells. (A) Overview of the six functional modules in hPGCdb. To support visualization and exploration, a user-friendly web interface for hPGCdb has been developed where users can browse, search, analyze, and download analytical results of interest. (B) A screenshot of the hPGCdb website homepage. (C) The genome browser tracks embedded in hPGCdb showing the ChIP-seq peaks around the gene POU5F1. (D-E) The 2D UMAP embedded in hPGCdb showing the cell populations of day49-112 ovary samples, and the corresponding expression of DDX4. Related to Figure 1.

**Figure S2.**
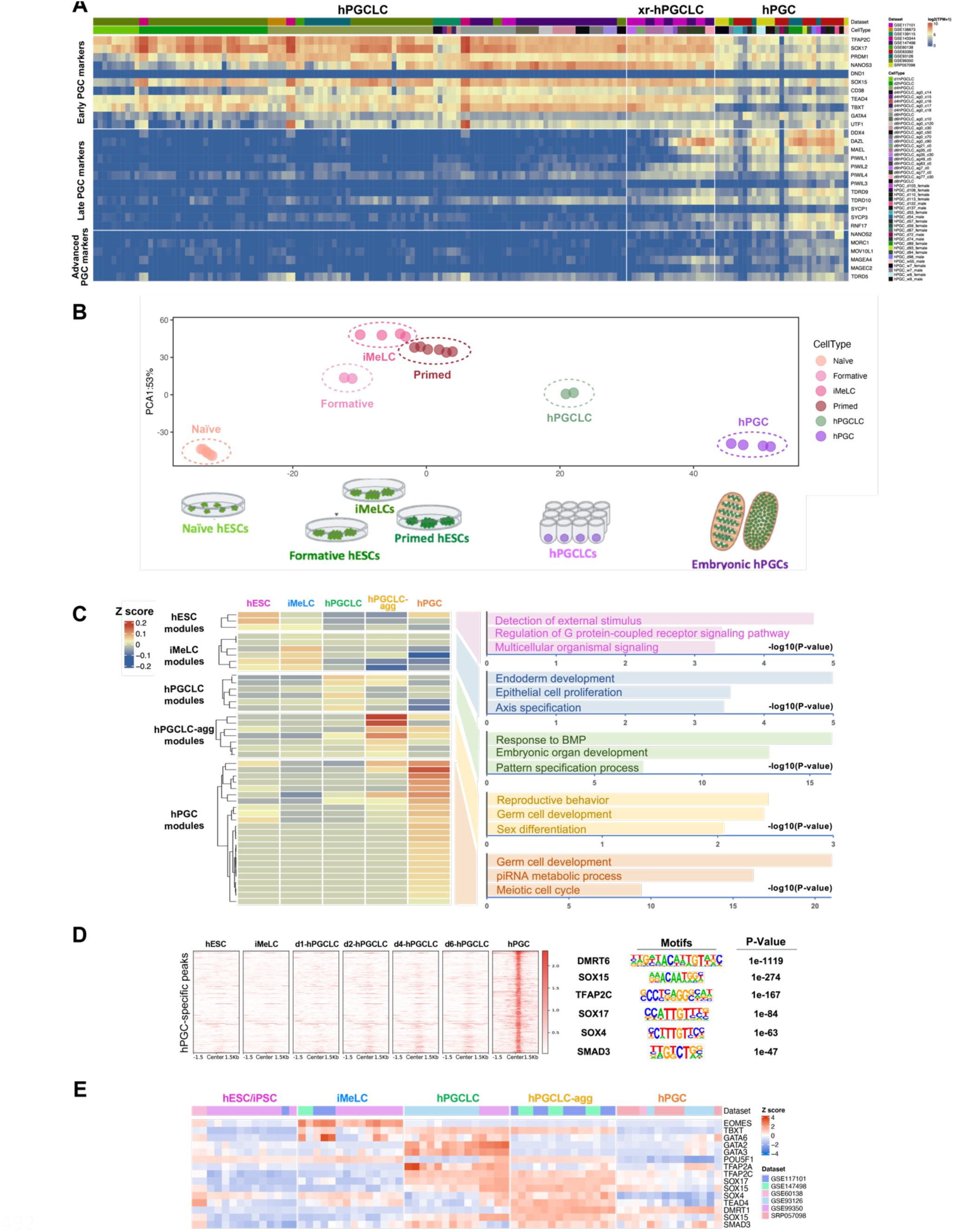
Analysis of the bulk RNA-seq data of human germ cells. (A) Heatmap showing the expression of the known germ cell marker at different stages. (B) PCA plot showing the developmental trajectory of hESCs from different pluripotent states (Naïve; Formative; Primed), iMeLCs, day4 hPGCLCs, and hPGCs. (C) Heatmap showing the average expression of gene modules at different stage of germline cells (left). Gene modules were clustered and labeled as stage-enriched modules for the five cell stages according to the average expression levels. The representative GO terms of each stage-enriched gene module group are displayed (right). (D) Heatmap showing the hPGC-specific ATAC-seq signals in hESCs, iMeLCs, day1 hPGCLCs, day2 hPGCLCs, day4 hPGCLCs, day6 hPGCLCs, and hPGCs, together with the corresponding TF motifs significantly enriched in these regions. (E) Heatmap showing gene expression level of candidate TF regulators across cell types, including hPSCs (hESCs/iPSCs), iMeLCs, hPGCLCs, xr-hPGCLCs, and hPGCs. Related to Figure 2.

**Figure S3.**
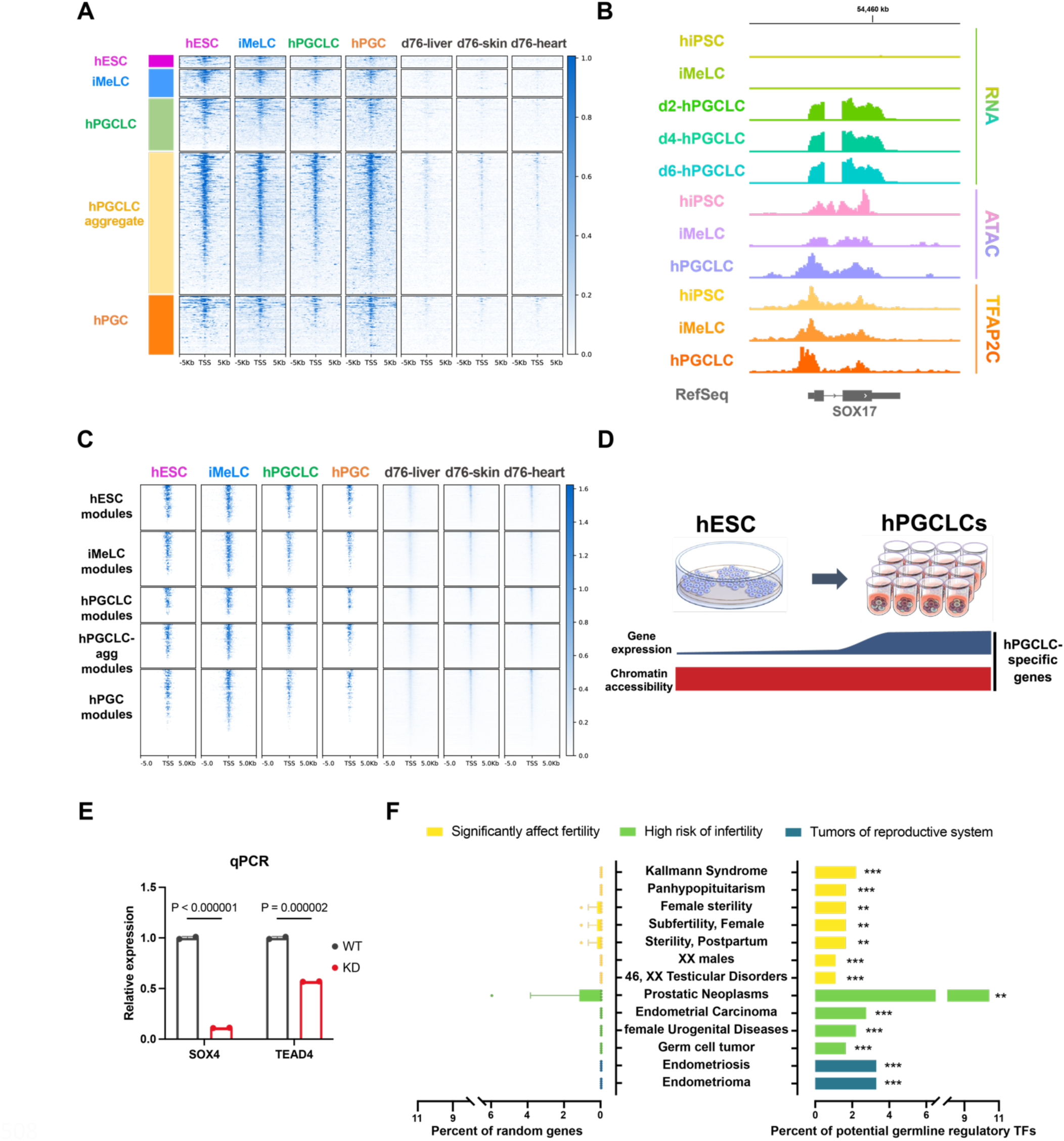
Analysis of putative transcription factors important for germline. (A) Metaplot showing the accessibility of chromatin regions around the stage-specific genes mentioned in Figue 2B for hESCs, iMeLCs, hPGCLCs, hPGCs, d76 liver, d76 skin, and d76 heart. (B) Genome browser tracks showing the gene expression around SOX17, the chromatin accessibility around SOX17, and the binding of TFAP2C to the genomic regions around SOX17. (C) Metaplot showing the accessibility of chromatin around the stage-enriched gene module group genes mentioned in Figure S2C for hESCs, iMeLCs, hPGCLCs, hPGCs, d76 liver, d76 skin, and d76 heart. (D) Schematic illustration showing that the germ cell genes are primed at the hESC stage. (E) Histogram showing the relative gene expressions of SOX4 and TEAD4 in control hESCs and knockdown hESCs assessed by qPCR. The average of two replicates are shown. Error bar, standard error of the mean (SEM). The statistical significance of the differences between Control and KD are evaluated by unpaired two-sided t-test assuming unequal variances. (F) Right panel shows histograms indicating the relationship between fertility diseases and the candidate TF regulators identified in Figure 3D. Left panel shows the corresponding random results (n = 5). Error bar, standard error of the mean (SEM). The statistical significance of the differences between random results and TF regulators’ results are evaluated by one sample t-test. Related to Figure 2.

**Figure S4.**
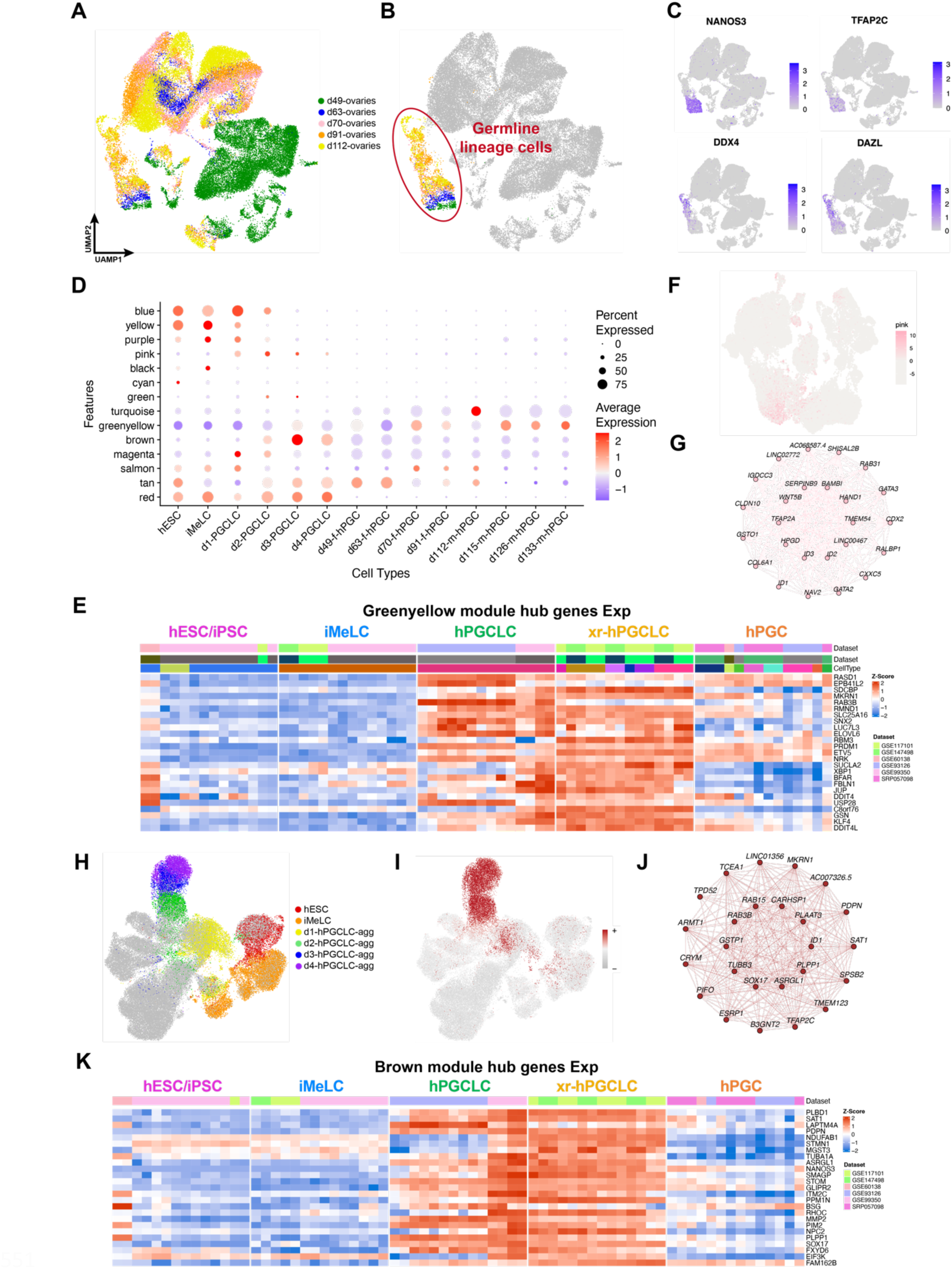
scWGCNA for identification of putative regulatory network modules for human germ cell fate. (A-B) UMAP plot showing all cells of day49-112 ovaries from the GSE143380 dataset. (C) UMAP plots showing the expression of two early germ cell marker genes (TFAP2C and NANOS3) as well as two late germ cell marker genes (DDX4 and DAZL) in day49-112 ovaries. (D) Dot plot showing the enrichment of different gene modules at different stages based on scWGCNA using the integrated germline lineage data. (E) The expression of hub genes identified from the germ cell regulatory network in Figure 3K. (F) UMAP plot showing the expression pattern of the putative regulatory network for TFAP2A+ progenitors based on scWGCNA. (G) The putative regulatory network for TFAP2A+ progenitors based on scWGCNA. (H) UMAP plot colored by the germline trajectory from scRNA-seq data of UCLA2 hESCs, iMeLCs, and day 1 to day 4 hPGCLCs. (I) UMAP plot showing the putative germ cell regulatory network based on scWGCNA using GSE140021-UCLA2 single cells. (J) The putative regulatory network module for germ cells based on scWGCNA using GSE140021-UCLA2 single cells. (K) The expression of genes identified from the germ cell regulatory network in Figure S4J. Related to Figure 3.

**Figure S5.**
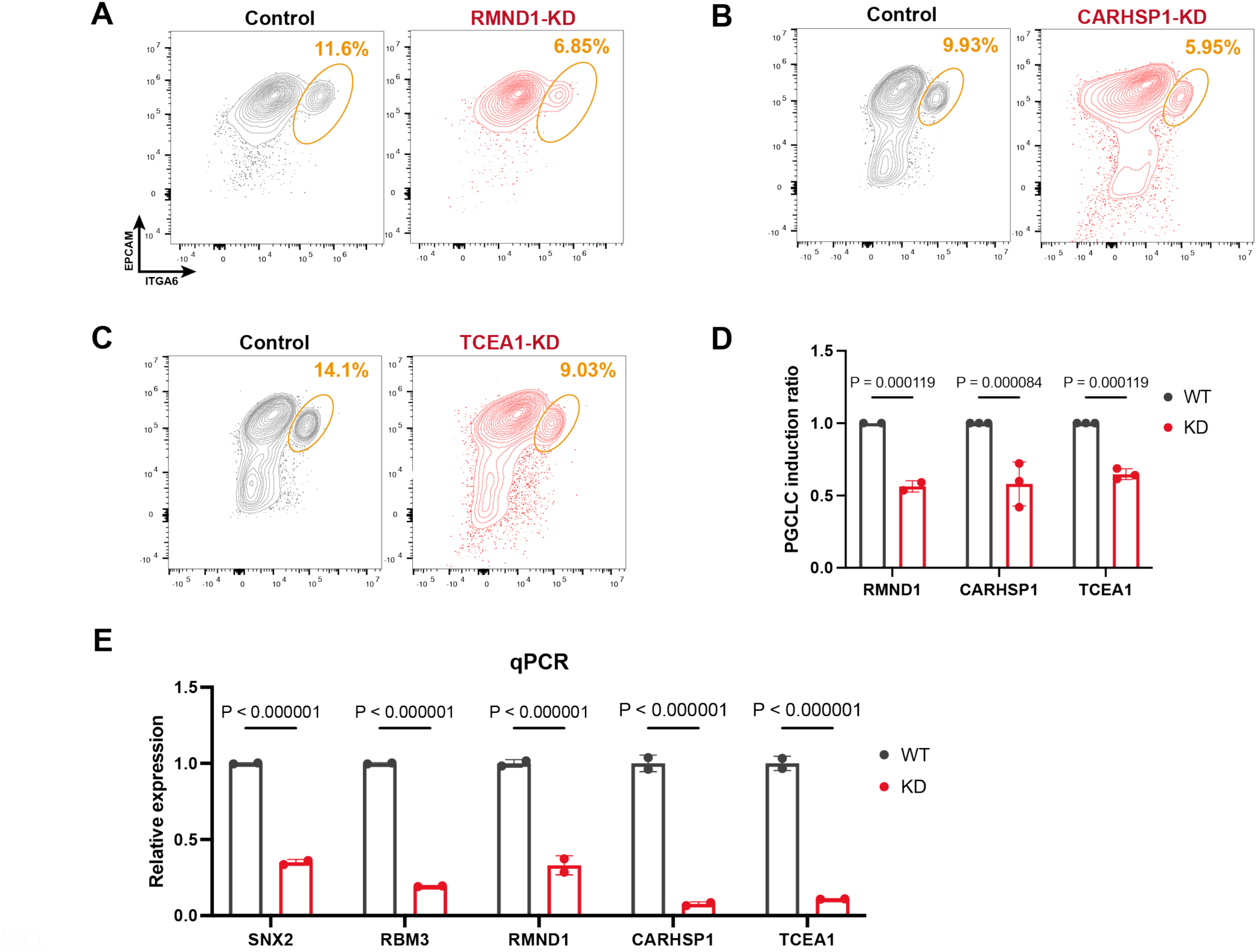
Examination of hPGCLC induction from knocking down of RMND1, CARHSP1, and TECA1. (A-C) Flow cytometry showing the induction of hPGCLCs at day 4 of aggregation differentiation using the control hESCs and CRISPRi-KD hESCs (A: RMND1-KD; B: CARHSP1-KD; C: TECA1-KD). (D) Histogram showing the scaled proportions of hPGCLCs derived from control hESCs and knockdown hESCs (RMND1-KD, TCEA1-KD, CARHSP1-KD) as assessed by flow cytometry. The average of two replicates are shown. Error bar, standard error of the mean (SEM). The statistical significance of the differences between Control and KD are evaluated by unpaired two-sided t-test assuming unequal variances. (E) Histogram showing the relative gene expressions of SNX2, RBM3, RMND1, CARHSP1, and TCEA1 in control hESCs and gene knockdown hESCs as assessed by qPCR. The average of two replicates are shown. Error bar, standard error of the mean (SEM). The statistical significance of the differences between Control and KD are evaluated by unpaired two-sided t-test assuming unequal variances. Related to Figure 3.

**Figure S6.**
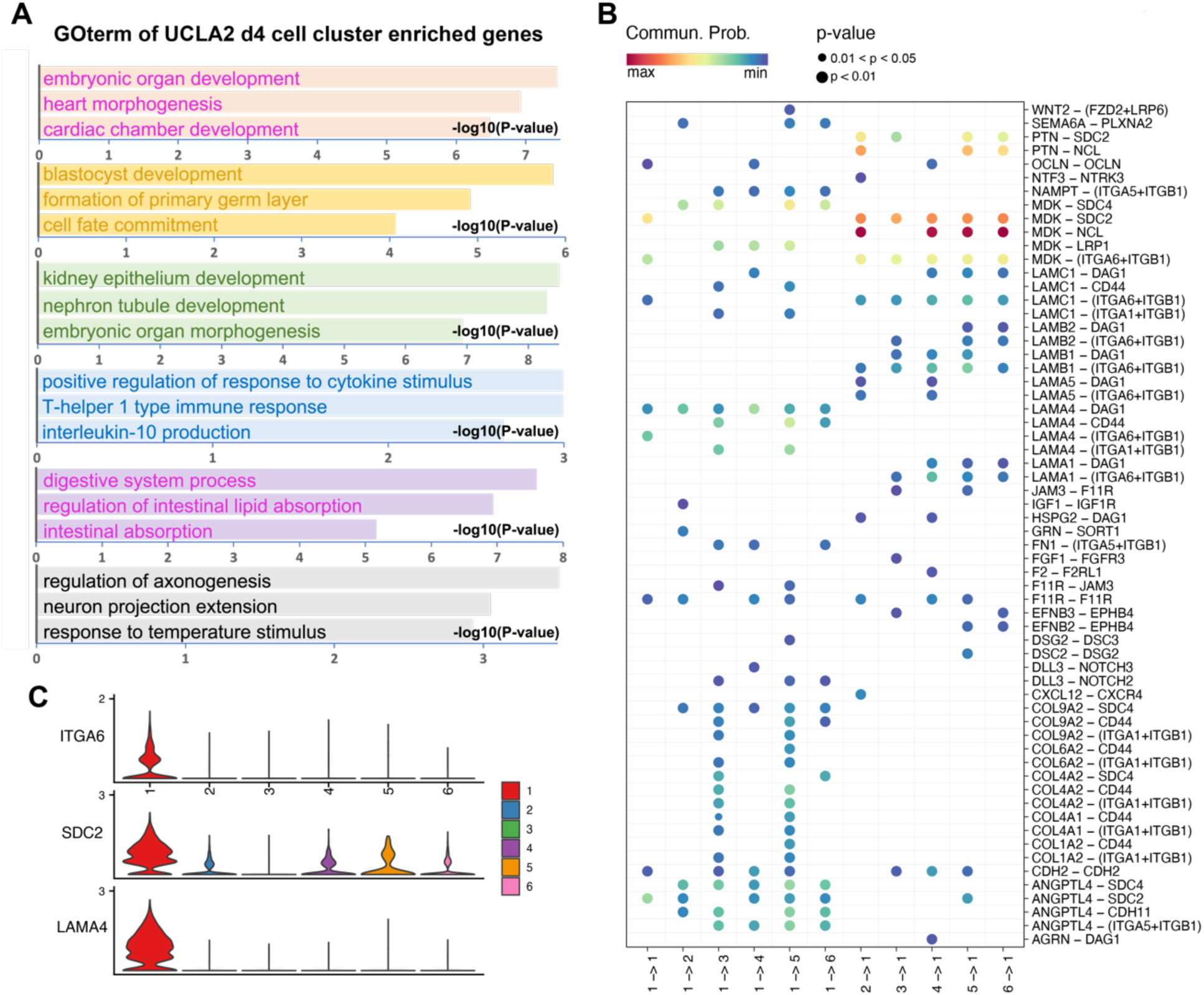
Elucidation of soma-germline interaction in hPGCLC aggregates. (A) GO terms enriched from cluster-specific genes of each cell cluster of UCLA2 day4 hPGCLC aggregates. The cell type of somatic cell clusters in UCLA2 day4 hPGCLC aggregates are annotated according to these GO terms. (B) Dot plot showing UCLA2 cell-cell interaction profile, including from day4 hPGCLC to soma and from soma to day4 hPGCLC. (C) Violin plot showing the expression profile of ITGA6, SDC2, and LAMA4 in all six cell clusters in UCLA2 day4 hPGCLC aggregates. Related to Figure 4.

**Figure S7.**
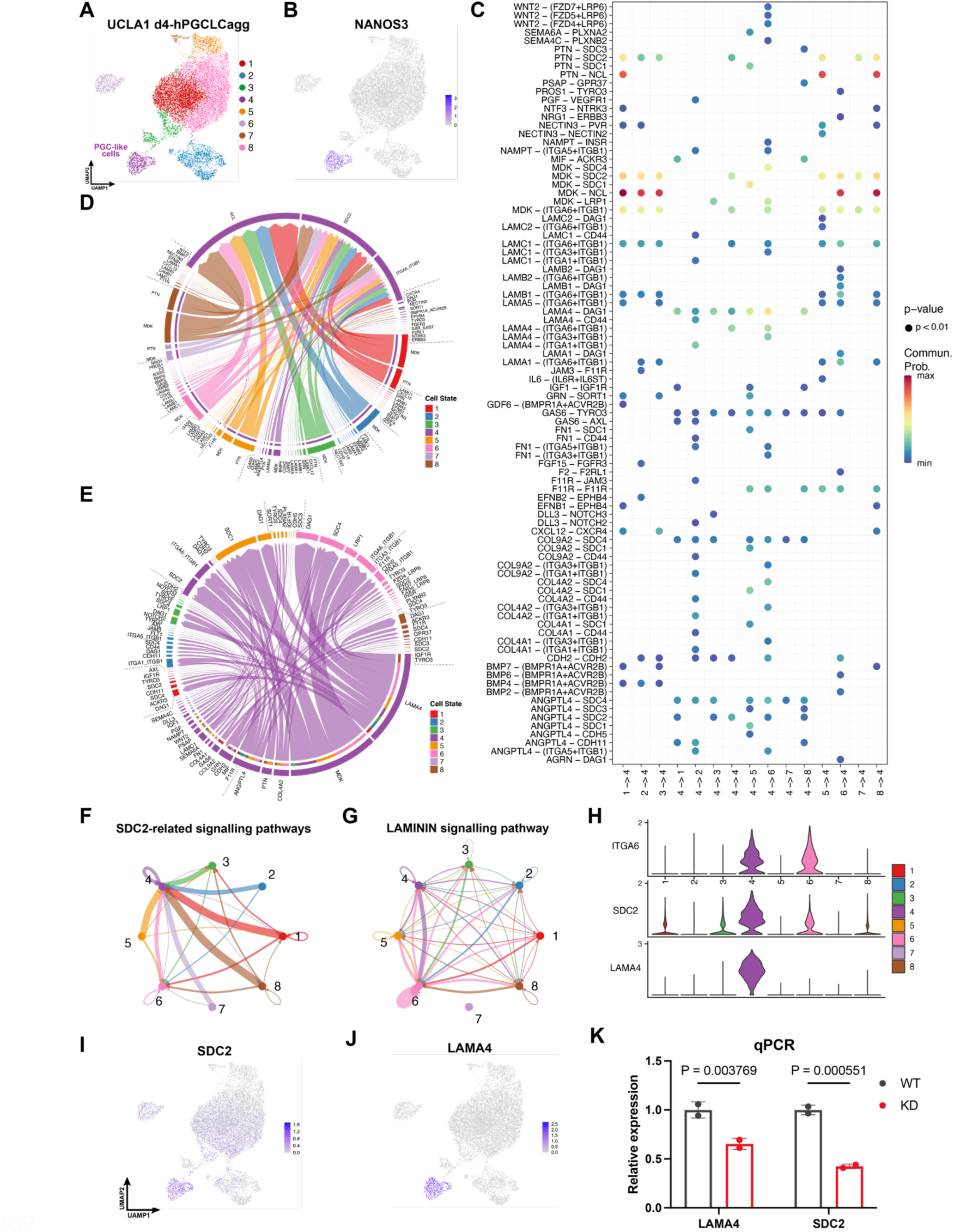
Elucidation of soma-germline interaction in UCLA1 hPGCLC aggregates. (A) UMAP plot showing the cells of UCLA1 day 4 hPGCLC aggregate samples. (B) UMAP plot showing the expression of NANOS3 in UCLA1 day 4 hPGCLC aggregate samples. (C) Dot plot showing UCLA1 cell-cell interaction profile, including from day4 hPGCLCs to soma and from soma to day4 hPGCLCs. (D) Chord plot showing cell-cell interactions targeting to UCLA1 day4 hPGCLCs from various somatic cell clusters. (E) Chord plot showing cell-cell interactions sourcing from UCLA1 day4 hPGCLC to various somatic cell clusters. (F) SDC2-related interactions among all cell clusters in UCLA1 day 4 hPGCLC aggregates. (G) LAMININ signaling-related interactions among all cell clusters in UCLA1 day 4 hPGCLC aggregates. (H) Violin plot showing the expression profile of ITGA6, SDC2, and LAMA4 in all UCLA1 eight cell clusters (I-J) UMAP plots showing the expression of SDC2 (I), LAMA4 (J) in UCLA1 day 4 hPGCLC aggregates, respectively. (K) Histogram showing the relative gene expressions of LAMA4 and SDC2 in control hESCs and knockdown hESCs assessed by qPCR. The average of three replicates are shown. Error bar, standard error of the mean (SEM). The statistical significance of the differences between control and KD are evaluated by unpaired two-sided t-test assuming unequal variances. Related to Figure 4.

**Figure S8.**
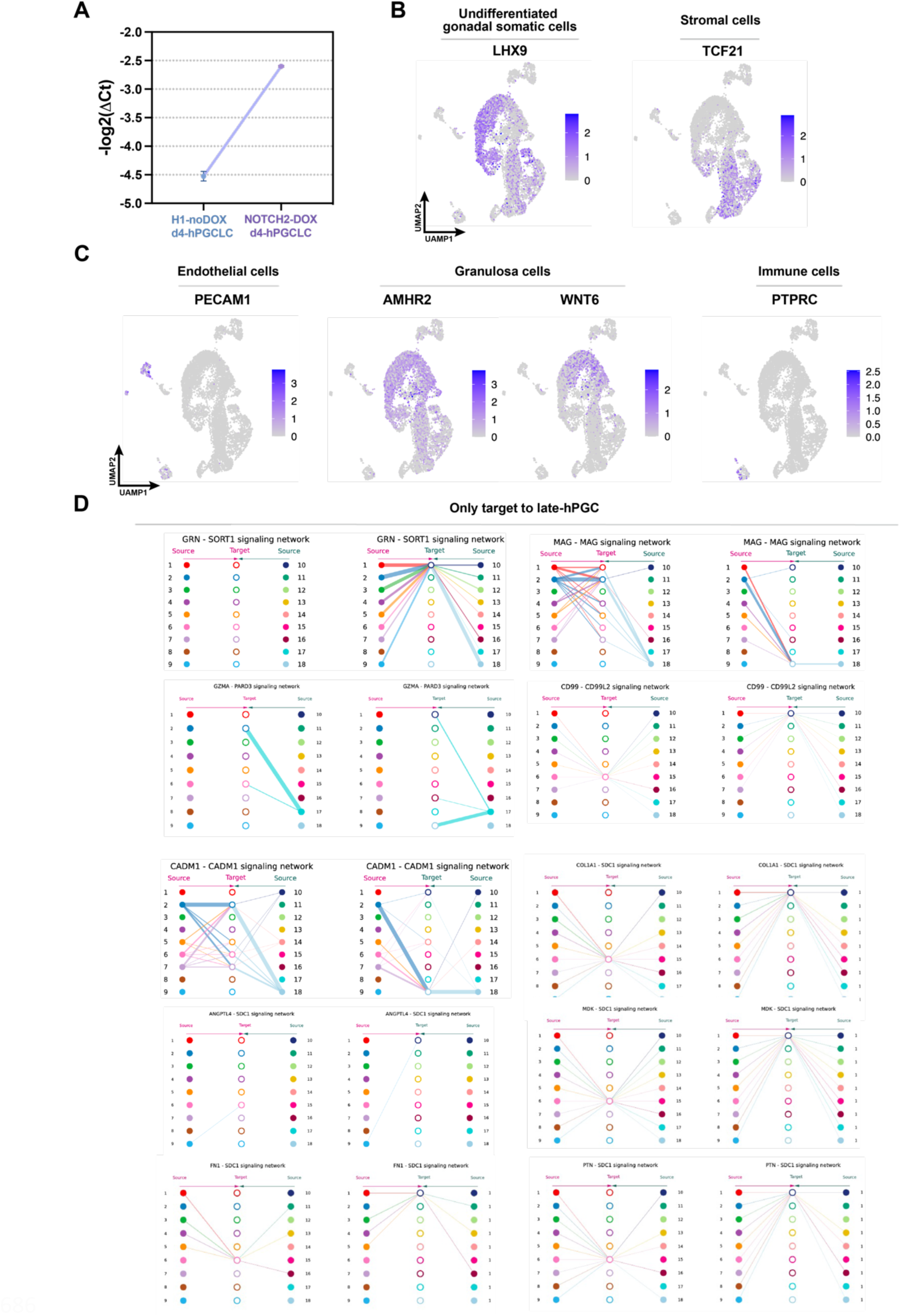
Analysis of soma-germline interaction for late PGCs. (A) Histogram showing NOTCH2 expression levels in control hESCs and NOTCH2-OE hESCs. The average of two replicates are shown. Error bar, standard error of the mean (SEM). The statistical significance of the differences between control and KD are evaluated by unpaired two-sided t-test assuming unequal variances. (B-C) The expression levels of the representative somatic marker genes for annotating somatic cell populations in Figure 5A. (D) The communication networks of the signaling from somatic cells to late PGCs. Related to Figure 5.

**Figure S9.**
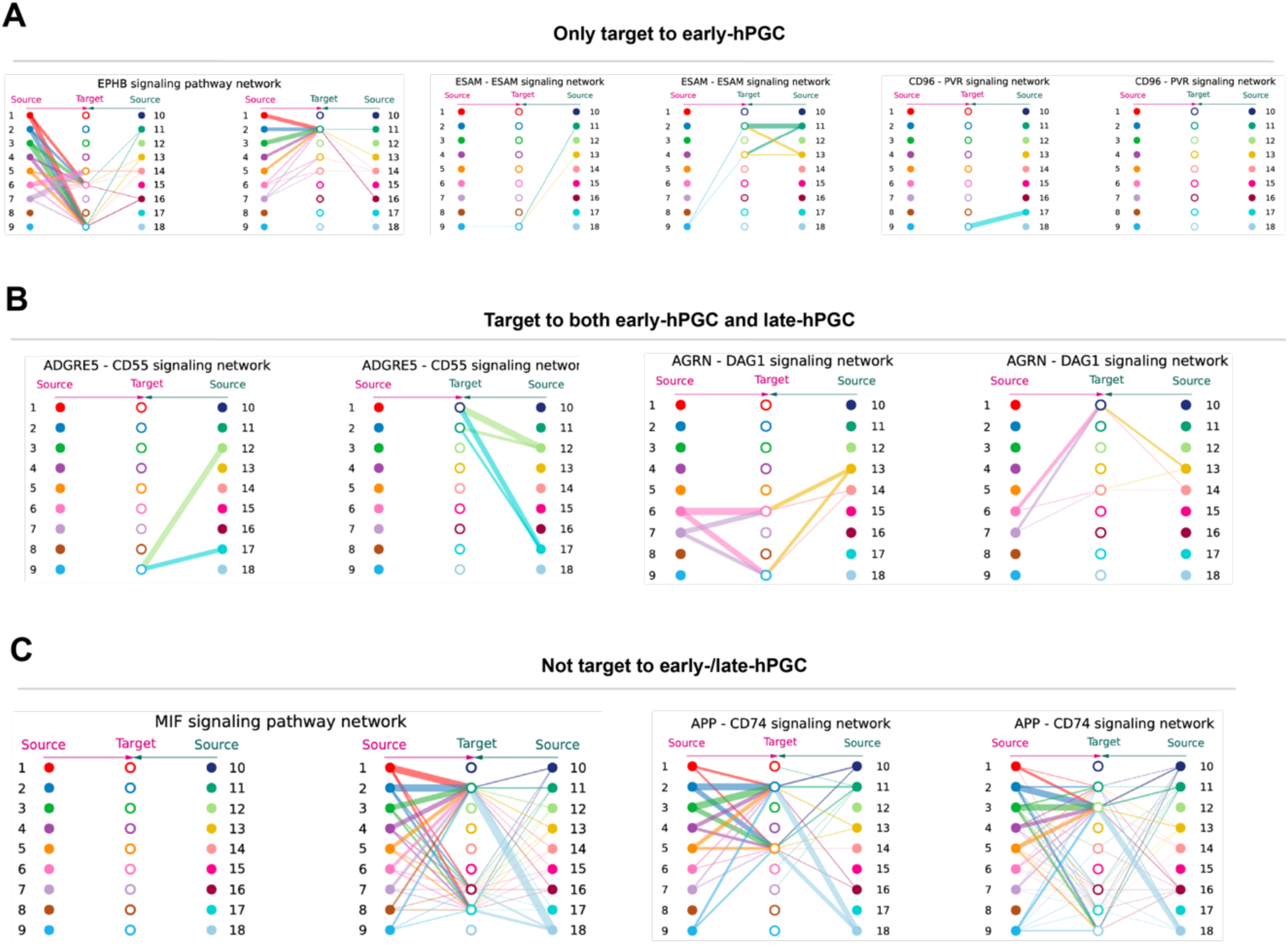
Soma-germline interaction examples for different kinds of signaling pathways. (A) The signaling pathways targeting to early PGCs but not late PGCs. (B) The signaling pathways targeting to both early PGCs and late PGCs. (C) The signaling pathways not targeting to either early PGCs or late PGCs. Related to Figure 5.

## MATERIALS AND METHODS

### DATA COLLECTION AND PROCESSING

hPGCdb was established from human PGC-related high-throughput sequencing datasets from GEO (https://www.ncbi.nlm.nih.gov/geo/) (1) and ArrayExpress (https://www.ebi.ac.uk/arrayexpress/) (2). A total of 683 samples across 27 Bulk RNA-seq, ATAC-seq, and ChIP-seq datasets, together with over three hundred thousand single cells in four scRNA-seq datasets, were collected and processed. A uniform analysis framework started by raw data was performed for all datasets. All published data used in this study is listed in Table S1.

### BulkRNA-seq data analysis

After checking the quality of each BulkRNA-seq sample by FastQC (https://github.com/s-andrews/FastQC), the quality and adapter trimming of raw reads were first performed using Trim-Galore (https://github.com/FelixKrueger/TrimGalore) and Cutadapt (https://github.com/marcelm/cutadapt). Reads were then aligned to hg38 using STAR (https://github.com/alexdobin/STAR) by allowing maximal two mismatches per read. Read counts per gene were calculated using FeatureCounts (https://subread.sourceforge.net/featureCounts.html).

Gene expression levels were normalized by both TPM (transcripts per kilobase million) and RPKM (reads per kilobase of exons per million aligned reads), and log2(TPM+1) and log2(RPKM+1) value were used for heatmap visualization. To integrate all Wide-Type samples, LIMMA (https://bioconductor.org/packages/release/bioc/html/limma.html) was used to remove batch effects between different datasets. After the integration, we used DESeq2 (https://bioconductor.org/packages/release/bioc/html/DESeq2.html) to perform Principal Component Analysis using the top 1000 variable genes.

Based on the expression levels of known gene markers of the germline cycle, seventy samples of the main five germline cell types across six datasets were selected for the identification of stage-specific genes. Limma and DESeq2 were used to remove batch effect and calculate significantly up-regulated genes at each cell type, respectively. Gene Ontology Enrichment analysis of the stage-specific genes was performed using clusterProfiler (https://github.com/YuLab-SMU/clusterProfiler).

### ATAC-seq and ChIP-seq data analysis

The quality and adapter control of ATAC-seq and ChIP-seq raw reads were the same as BulkRNA-seq data. Reads were then aligned to hg38 using bowtie (https://github.com/BenLangmead/bowtie) or bowtie2 (https://github.com/BenLangmead/bowtie2) depending on the read length. After the PCR duplicate removal by Sambamba (https://github.com/biod/sambamba), the reads with Phred Quality Score < 30 or mapped to mitochondrial genes were discarded. Besides, paired-end reads mapped in improper pair were discarded as well. ATAC-seq and ChIP-seq signals were then normalized as TPM to display corresponding tracks on the hPGCdb website.

To identify and visualize stage-specific peaks of day1-day4 hPGCLCs, DiffBind (https://bioconductor.org/packages/release/bioc/html/DiffBind.html) was used with p-value < 0.005. For the identification of enriched motifs in different stage-specific peak sets, findMotifs.pl function in HOMER (http://homer.ucsd.edu/homer/) was utilized to find both known motifs and 12-mer *de-novo* motifs. Metaplots were visualized with deeptools (https://github.com/deeptools/deepTools), and the ATAC-seq peak enrichment score analysis was performed using cisDynet(https://tzhu-bio.github.io/cisDynet_bookdown/).

### scRNA-seq data analysis

#### Pre-processing

All the four single cell RNA-seq datasets are from 10X Genomics platform. Cell Ranger with its built-in reference *refdata-gex-GRCh38-2020-A.tar.gz* was used to construct UMI (unique molecular identifier) count matrices for each dataset. Afterwards, Scrublet (https://github.com/swolock/scrublet) was applied to remove doublets (4% per 10,000 cells), and Seurat R package (https://satijalab.org/seurat/) was then utilized to perform a series of analyses: cell quality control (200 ≤ nfeatures ≤ 2500 and MT < 20%), gene filtration (min.cells > 3), expression normalization (log-transeformed and multipled by 10,000), expression scaling with variable genes, PCA (Principal Component Analysis) dimension reduction, and UMAP (Uniform Manifold Approximation and Projection) clustering using the top 50 principal components and min_dist = 0.9.

Inside each dataset, batch correction between biological replicates were performed using functions *FindIntegrationAnchors* and *IntegrateData* with default parameters. Seurat’s canonical correlation analysis procedure was also utilized for the datasets containing multi-batches. The top 30 canonical correlation vectors were saved for the following germline lineage analysis.

#### Germline lineage extraction and analysis

For one-batch datasets and multi-batch dataset, the scaled expression matrices and the top 30 CCA components matrices were used to build the lineage tree by tcURD (https://github.com/farrellja/URD), respectively. Based on tcURD tree and the expressions of the early PGC marker, NANOS3, as well as the late PGC marker, DDX4, the germline lineage and corresponding cells of each dataset were determined and extracted with pseudo-time information. The extracted germline lineage cells from each dataset were merged to generate an integrated germline lineage object, where the batch effect in different studies were corrected by MNN (Mutual Nearest Neighbors) method. The integrated germline lineage object was then performed UMAP clustering and pseudo-time analysis using Monocle3 (https://cole-trapnell-lab.github.io/monocle3/). A group of germline gene markers were investigated to verify the validation of the data integration and pseudo-time results.

The single-cell Weighted Correlation Network Analysis (scWGCNA) was also applied to both the integrated germline lineage dataset and the GSE140021 UCLA2 dataset to identify unbiased gene modules using R package scWGCNA (https://github.com/CFeregrino/scWGCNA). Beforehand, Seurat *FindIntegrationAnchors*, *IntegrateData, and NormalizeData* were carried out to generate a batch-effect-removed and normalized expression matrix as the input of scWGCNA.

#### Day4 hPGCLC aggregate and week10 female ovary analysis

COSG (https://github.com/genecell/COSG) was used to identify cluster-enriched genes with higher cell type specificity. Gene Ontology Enrichment analysis of cell cluster-enriched genes were performed to annotate each somatic cell cluster. Then CellChat (https://github.com/sqjin/CellChat) was utilized to identify legend-receptor pairs with important roles in d4 hPGCLC aggregates.

The UCLA1/2 day4 hPGCLC aggregates samples and the week 10 ovary samples were firstly pre-processed as mentioned before, including batch effect removal, normalization, scaling, PCA analysis, and neighbor determination. Afterwards, clustering of the day4 hPGCLC samples and the week 10 ovary sample were determined with resolution = 0.2 and resolution = 0.6, respectively.

UMAP of both two kind of samples were then calculated by RunUMAP function in Seurat package using top50 PCs and min_dist = 0.75.

For somatic cell cluster annotation in d4hPGCLC aggregates, COSG was first used to identify cluster-enriched genes with high cell type specificity. Gene Ontology Enrichment analysis of somatic cell cluster-enriched genes was performed to annotate each somatic cell cluster.

The somatic cell cluster in week10 female ovary samples were annotated using female soma marker from 2022_Min_Chen.

Then, in d4 hPGCLC aggregates, CellChat was utilized to identify legend-receptor pairs with important roles, while in week10 female ovary samples, CellChat was utilized to analyze the intercellular communication between somatic cells and hPGCLCs/hPGCs.

## DATABASE IMPLEMENTATION

The hpGCdb database was deployed on the Tomcat web server and developed using Spring Boot as the basic architecture of the database website. Bootstrap, thymeleaf, HTML5, CSS and JQuery libraries were used to set up the user interfaces. MySQL was applied to perform data management and Mybatis was used to access the database. Java script plugins such as Echarts.js and plotly.js were applied to build interactive plots on the website. Datatables.js was used to view data information and igv.js served as interactive genome visualization tool. The bioinformatic analyses were done by the R framework and python scripts. Users can quickly locate sequencing data of interest, such as RNA-seq, ATAC-seq, ChIP-seq and scRNA-seq to use the corresponding online analysis tools provided.

## EXPERIMENTS

### gRNA plasmid construction

Plenti-dual gRNA plasmid backbone (a gift from Chan’s Lab) was digested with BbSI. Two gRNAs were designed within primers, the PCR product of which was ligated into the backbone through Gibson assembly to achieve gRNA cloning. Sanger sequencing was performed to check the sequence of inserted gRNAs.

### Lentiviral production

293T cells in 6-well plates were transfected at 70-80% confluency. 3200ng plenti-dual gRNA plasmid, 800 ng pMD2.G and 2400 ng psPAX2 were delivered into cells by Lipofectamine 2000.

72h after transfection, lentiviral supernatant was collected and filtered through 0.45mm filter. Lentivirus was then concentrated by addition of PEG8000 and centrifuge at 4000g, followed by resuspension in 200ul ddH2O.

### CRISPR knock-down

pC13N-dCas9-BFP-KRAB, CLYBL-L, CLYBL-R constructs were transfected into the hESCs with P3 Primary Cell 4D Nucleofector X Kit S (Lonza) on 4D-nucloefector (program CA137, Lonza) following manufacture’s protocol. 8*105 hESCs and 4μg of each plasmid were used for nucleofection. BFP positive cells were sorted and continuously cultured to reach 100% positive rate. Lentivirus of targeted dual gRNA plasmid were transduced into this cell line to achieve knockdown.

### hESC culture and hPGCLC induction

hESCs were maintained on mouse embryonic fibroblasts (MEFs) in hESC media (20% knockout serum replacement, 100mM L-Glutamine, 1× MEM Non-Essential Amino Acids, 55mM 2-Mercaptoethanol, 10ng/mL recombinant human FGF basic, 1× Penicillin-Streptomycin, and 50ng/mL primocin, DMEM/F12 media). hESCs were dissociated into single cells with Accutase for 5min, and single cell suspension of 300k hESCs were plated on Human Plasma Fibronectin (HPF)-coated 12-well-plates, cultured in 2mL iMeLC media (15% KSR, 1× MEM Non-Essential Amino Acids, 0.1mM 2-Mercaptoethanol, 1× Penicillin-Streptomycin-Glutamine, 1mM sodium pyruvate, 50ng/mL Activin A, 3mM CHIR99021, 10mM of ROCKi (Y27632), GMEM). After 24 hours, iMeLCs were dissociated into single cells with Accutase for 5min, and single cell suspension of 40k iMeLCs were plated on Anti-Adherence Rinsing Solution-coated U-bottom 96-well-plates, cultured in 200μL PGCLC media/well (day 0) (15% KSR, 1× MEM Non-Essential Amino Acids, 0.1mM 2-Mercaptoethanol, 1× Penicillin-Streptomycin-Glutamine, 1mM sodium pyruvate, 10ng/mL human LIF, 200ng/mL hu-man BMP4, 50ng/mL human EGF, 10mM of ROCKi (Y27632) and GMEM). hPGCLC aggregates were collected for analysis at day4.

### Flow cytometry

hPGCLC aggregates were dissociated into single cells by 0.05% trypsin for 10min. The harvested cells were stained with 7-AAD viability dye and antibodies including ITGA6 conjugated with BV421 (BioLegend, 313624, 1:200), EPCAM conjugated with 488 (BioLegend, 324210, 1:200), EPCAM conjugated with APC (BioLegend, 324207, 1:200). The signals were detected by flow cytometry using Flow Cytometer (Cytek Biosciences). hPGCLCs was identified as ITGA6 and EPCAM double positive population.

### qRT-PCR

RNA weas extracted using FastPure Cell/Tissue Total RNA Isolation Kit (Vazyme, Cat#RC101). Reverse transcription was performed following the instructions of Evo M-MLV RT Premix (Accurate biology, Cat#AG11706). qRT-PCR was performed following the instruction of SYBR® Green Premix Pro Taq HS qPCR Kit (Accurate biology, Cat#AG11701) in Bio-Rad CFX96 PCR machine. The qRT-PCR data were normalized to the expression of housekeeping gene GAPDH. The primers for qRT-PCR are listed below:

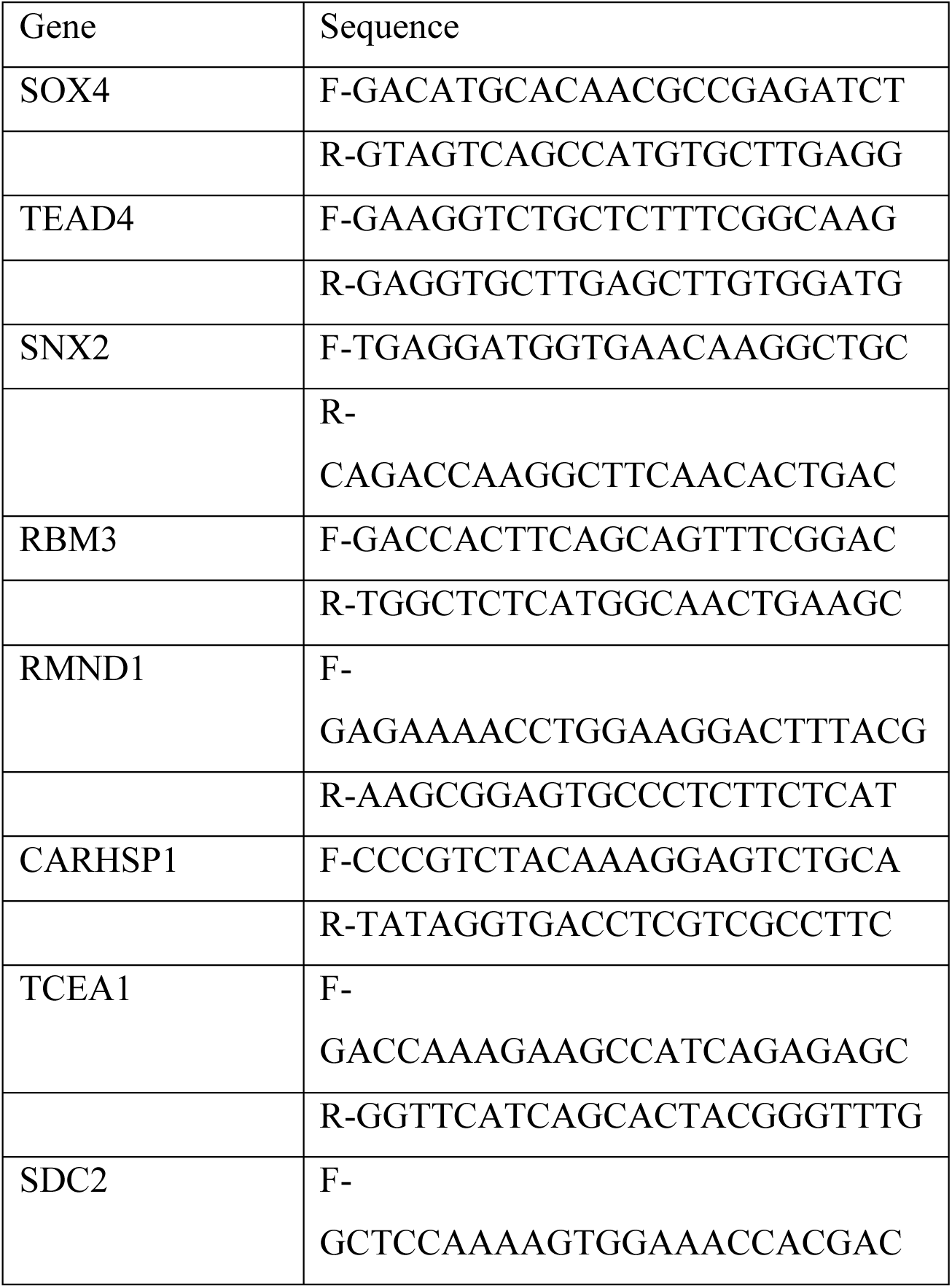
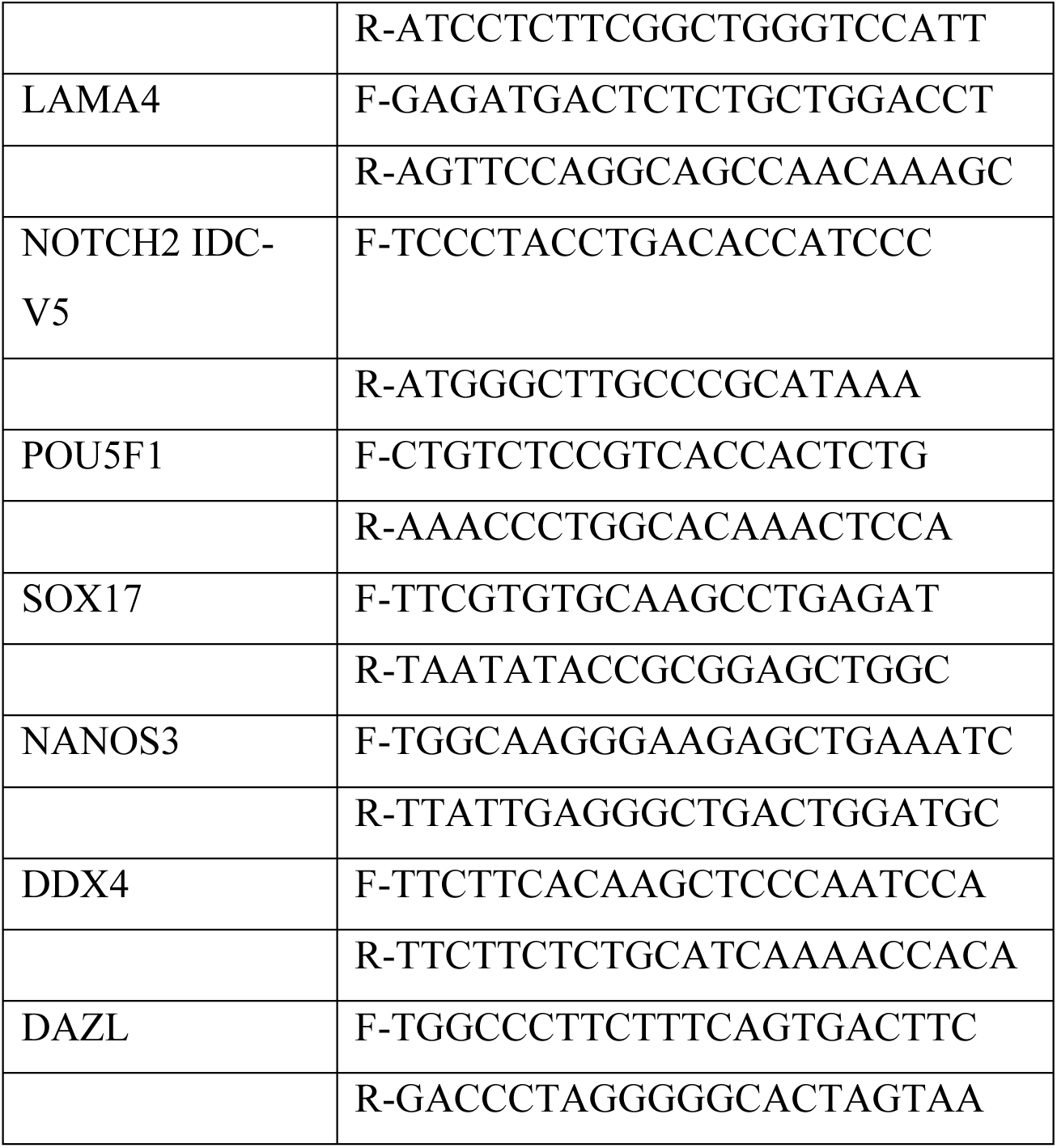

